# Single-molecule identification of the target RNAs of different RNA binding proteins simultaneously in cells

**DOI:** 10.1101/2022.09.06.506842

**Authors:** Mathieu N. Flamand, Ke Ke, Renee Tamming, Kate D. Meyer

## Abstract

RNA-binding proteins (RBPs) regulate nearly every aspect of mRNA processing and are important regulators of gene expression in cells. However, current methods for transcriptome-wide identification of RBP targets are limited since they examine only a single RBP at a time and since they do not provide information on the individual RNA molecules that are bound by a given RBP. Here, we overcome these limitations by developing TRIBE-STAMP, an approach for single-molecule detection of the target RNAs of two RNA binding proteins simultaneously in cells. We apply TRIBE-STAMP to the cytoplasmic m^6^A reader proteins YTHDF1, 2, and 3 and discover that individual mRNA molecules can be bound by more than one YTHDF protein throughout their lifetime, providing new insights into the function of YTHDF proteins in cells. TRIBE-STAMP is a highly versatile approach that enables single-molecule analysis of the targets of RBP pairs simultaneously in the same cells.

## Introduction

Identifying the RNA targets of RBPs in cells has been important for our understanding of how individual RBPs control gene expression. To achieve this, methods such as RIP-seq or CLIP-seq and its variants have often been employed. However, these strategies are limited to examining a single RBP at a time, which makes it difficult to study potential synergistic or competitive RNA targeting between two distinct RBPs in the same cells. Moreover, since immunoprecipitated RNA fragments are typically subjected to bulk sequencing, RIP/CLIP-based approaches do not provide information on individual RNA molecules that are bound by a given RBP and instead only identify RNAs at the gene level.

Recently, TRIBE/HyperTRIBE and STAMP have been developed as alternative methods for transcriptome-wide mapping of the RNA targets of RBPs (McMahon et al. 2016; Xu et al. 2018; Brannan et al. 2021). Both methods involve fusing an RBP to the catalytic domain of a deaminase enzyme (ADAR in the case of TRIBE; APOBEC1 in the case of STAMP). When these fusion proteins are expressed in cells, they direct A-to-I or C-to-U editing, respectively, at residues in proximity to RNA binding sites. These approaches have many advantages, including the ability to mark the individual RNA molecules that are bound by an RBP. However, to date, TRIBE and STAMP have been used in isolated studies focused on identifying the RNA targets of only a single RBP at a time.

We reasoned that we could take advantage of the unique mutation signatures induced by TRIBE and STAMP to simultaneously identify the mRNA targets of two RBPs at once. By co-expressing one RBP fused to ADAR and a different RBP fused to APOBEC1 in the same cells, we can examine each deamination event (A-to-I or C-to-U, henceforth referred to as A2I and C2U) and determine whether different RBPs bind to the same mRNA molecules or to distinct pools of mRNAs in cells. Using this method, which we call TRIBE-STAMP, we report here the ability to simultaneously detect the molecular RNA targets of pairs of distinct RBPs in cells.

To demonstrate the utility of the TRIBE-STAMP approach, we applied it to the YTHDF family of cytoplasmic m^6^A reader proteins. This includes three paralogs (YTHDF1, YTHDF2, and YTHDF3; hereafter DF1, DF2, and DF3) which share a C-terminal YTH domain responsible for m^6^A recognition. Although initially proposed to bind unique subsets of cellular mRNAs and carry out distinct functions, more recent models suggest that these three proteins function redundantly to promote mRNA decay (Wang et al. 2014; Wang et al. 2015; Li et al. 2017; Shi et al. 2017; Rauch et al. 2018; Shi et al. 2019; Kontur et al. 2020; Lasman et al. 2020; Zaccara and Jaffrey 2020). To address these discrepancies, we applied TRIBE-STAMP to each combination of DF proteins and found a high degree of overlap among the target mRNAs of all three proteins. We discovered that individual mRNA molecules can be bound by more than one DF protein throughout their lifetime, arguing against the idea that distinct transcripts are targeted uniquely by individual DF proteins. Furthermore, our data reveal sequential binding of distinct DF proteins to individual mRNA molecules and are consistent with a model in which DF1 and DF3 do not immediately promote mRNA decay. We anticipate that TRIBE-STAMP can be used across diverse combinations of RBPs to provide similar insights into other RNA:protein interactions and to deepen our understanding of RBP function in cells.

## RESULTS

### Simultaneous detection of the target mRNAs of distinct RBPs with TRIBE-STAMP

To determine whether TRIBE-STAMP can enable the simultaneous mapping of the targets of two individual RBPs in cells at the same time, we applied it to the cytoplasmic m^6^A reader proteins DF1, DF2, and DF3. We co-expressed DF-ADAR and DF-APOBEC1 fusion proteins in HEK293T cells by first generating stable cell lines expressing inducible APOBEC1-DF1,2, or 3 (Fig. 1A and Supplemental Fig. S1A,B). We then transfected each stable cell line with plasmids expressing the human ADAR2 catalytic domain (ADARcd-E488Q (Xu et al. 2018), henceforth referred to as ADAR) fused to DF1,2 or 3 and treated cells with doxycycline for 24h to induce DF-APOBEC1 expression. The stable cell lines and transfected plasmids contained EGFP and mCherry markers, respectively, enabling sorting of green/red fluorescent cells by flow cytometry to ensure that all analyzed cells expressed both the ADAR and APOBEC1 fusion proteins (Fig. 1a). This strategy enabled robust and concurrent expression of each combination of DF protein pairs (DF1+2, DF1+3, or DF2+3) (Supplemental Fig. S1C,D).

**Fig. 1:**
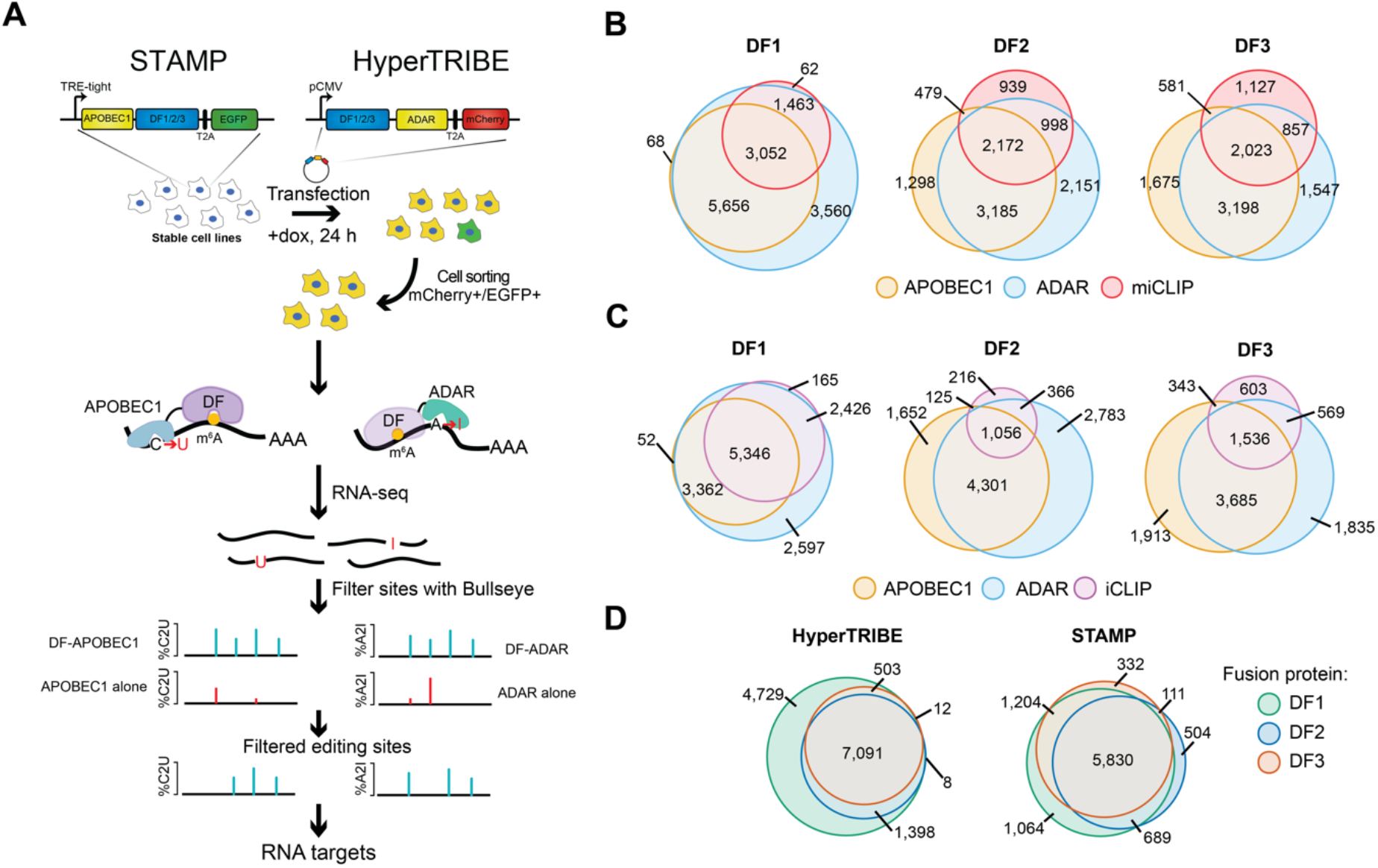
TRIBE-STAMP simultaneously identifies the target mRNAs of distinct DF proteins in cells. **A,** Overview of the TRIBE-STAMP approach used to identify DF protein targets. **B**, DF-ADAR and DF-APOBEC1 target mRNAs have a high degree of overlap. Euler plots show the overlap of target mRNAs identified by HyperTRIBE (DF-ADAR) or STAMP (DF-APOBEC1) fusions for each DF protein. **C**, m^6^A-containing mRNAs are edited by DF-ADAR and DF-APOBEC1 fusion proteins. Euler plots show the overlap between target mRNAs of DF-ADAR and DF-APOBEC1 fusion proteins and methylated mRNAs identified by miCLIP (Linder et al. 2015). **D**, DF target mRNAs identified by iCLIP are also identified by DF-ADAR and DF-APOBEC1 fusion proteins. Euler plots show the overlap between DF targets identified by DF-ADAR and DF-APOBEC1 fusion proteins and mRNAs containing DF iCLIP peaks (Patil et al. 2016).

We first sought to confirm that TRIBE-STAMP can be used to faithfully identify the RNA targets of individual DF proteins. We performed RNA-seq on cells co-expressing the ADAR and APOBEC1 fusion proteins for a particular DF protein. To identify A2I and C2U editing events transcriptome-wide, we developed a modified version of Bullseye, a pipeline we previously created to identify C2U editing events introduced by the m^6^A profiling method DART-seq(Meyer 2019; Flamand and Meyer 2022; Tegowski et al. 2022). Briefly, we optimized Bullseye for detection of both A2I and C2U mutations that are enriched relative to reference samples expressing the ADAR or APOBEC1 proteins alone. We then employed stringent filtering parameters to retain only the editing events which occur in multiple biological replicates and which are not caused by endogenous single nucleotide polymorphisms (SNPs) (see Methods). Using this approach, we identified thousands of target mRNAs for each fusion protein (Supplemental Table S1). Editing rates (%C2U or %A2I) at individual sites were similar across all three DF proteins when tethered to either ADAR or APOBEC1, suggesting that DF proteins do not differentially influence the efficiency of either deaminase enzyme (Supplemental Fig. S2A-C). Additionally, when we compared the mRNA targets identified by the ADAR and APOBEC1 versions of the same DF protein, we found that the majority of target mRNAs are bound by both fusion proteins, indicating that the ADAR and APOBEC1 fusions identify similar mRNAs for each DF protein (Supplemental Fig. S2D).

Consistent with the role of DF proteins as m^6^A readers, we observed a high degree of overlap between DF-ADAR and DF-APOBEC1 target mRNAs and methylated mRNAs identified by miCLIP (Linder et al. 2015) (Fig. 1B). Transcripts with more m^6^A sites also contain a larger number of editing sites for all DF fusion proteins (Supplemental Fig. S2E). Additionally, we found that mRNAs edited by DF-ADAR or DF-APOBEC1 overlapped well with DF target mRNAs identified by iCLIP (Patil et al. 2016), further confirming the accuracy of the TRIBE-STAMP approach (Fig. 1C and Supplemental Fig. S2F). Interestingly, we found that DF1 consistently targeted the most mRNAs and edited more sites per transcript compared to DF2 and DF3, regardless of whether it was fused to ADAR or APOBEC1 (Fig. 1D and Supplemental Fig. S2G). This was confirmed with analysis of endogenous DF iCLIP data (Patil et al. 2016), which also revealed a greater number of target mRNAs for DF1 compared to DF2 or DF3 in HEK293T cells (Fig. 1D and Supplemental Table S2). Altogether, these data demonstrate that tagging DF proteins with either ADAR or APOBEC1 enables the accurate identification of DF target mRNAs in cells.

### TRIBE-STAMP reveals RBP binding to individual RNA molecules

Previous studies have used CLIP-based methods to identify YTHDF-bound transcripts, but these methods provide only a snapshot of RNA:protein interactions and are limited to examining a single protein at a time. In contrast, TRIBE-STAMP allows simultaneous identification of the RNA targets of two proteins at once. We therefore examined the mRNA targets that are shared across the DF proteins by identifying the mRNAs that have both A2I and C2U editing in cells co-expressing each combination of DF fusion protein pairs. We found that each pair of DF proteins binds to largely the same mRNAs in cells, regardless of which tag (ADAR or APOBEC1) is used (Supplemental Fig. S2F). These data are consistent with iCLIP studies and suggest that DF proteins mostly recognize the same target mRNAs in cells (Lasman et al. 2020; Zaccara and Jaffrey 2020).

The finding that different DF proteins share most of the same target mRNAs has been used to support the hypothesis that DF proteins function redundantly to ensure a particular fate (destabilization) for methylated mRNAs (Kontur et al. 2020; Lasman et al. 2020; Zaccara and Jaffrey 2020). However, analysis of DF targets at the gene level does not discriminate between individual mRNA molecules. For instance, there may be many copies of a particular target mRNA in cells, and each DF protein may bind to its own unique subset of these mRNA molecules. Such a scenario would further support the functional redundancy model, with each mRNA molecule being bound by a single DF protein and then targeted for degradation. Conversely, if individual transcript copies are bound by multiple DF proteins throughout their lifetime, this would suggest that at least some DF proteins do not immediately initiate the degradation of their targets and instead potentially carry out additional functions. Because TRIBE-STAMP introduces irreversible mutation signatures into target transcripts, single-molecule binding information can be uncovered by assessing A2I and C2U edits that occur within individual sequencing reads which represent distinct mRNA molecules. Reads that contain both A2I and C2U edits would therefore reflect mRNA molecules bound by two distinct fusion proteins.

We analyzed TRIBE-STAMP data for each pair of co-expressed DF fusion proteins to determine whether individual reads contain both A2I and C2U edits. First, we considered all of the shared target mRNAs identified in cells co-expressing a given pair of DF proteins (Supplemental Fig. S2D). We then identified, within each set of common target mRNAs, pairs of A2I and C2U editing sites that are found within a 150 nt window on individual reads. The 150 nt distance was chosen based on the average read length of each dataset and ensures that there is the potential to capture co-editing events on a single read for that mRNA. Within this refined set of common target mRNAs for each pair of co-expressed DF proteins, we then identified the pairs of editing sites with evidence for co-editing (defined as having at least 2 reads with both A2I and C2U editing within the read) in all biological replicates (Fig. 2A, Supplemental Table S3 and Methods). This analysis enables identification of the proportion of shared target mRNAs for a given pair of DF proteins that show evidence for co-binding of the DF proteins to the same RNA molecules.

**Fig. 2:**
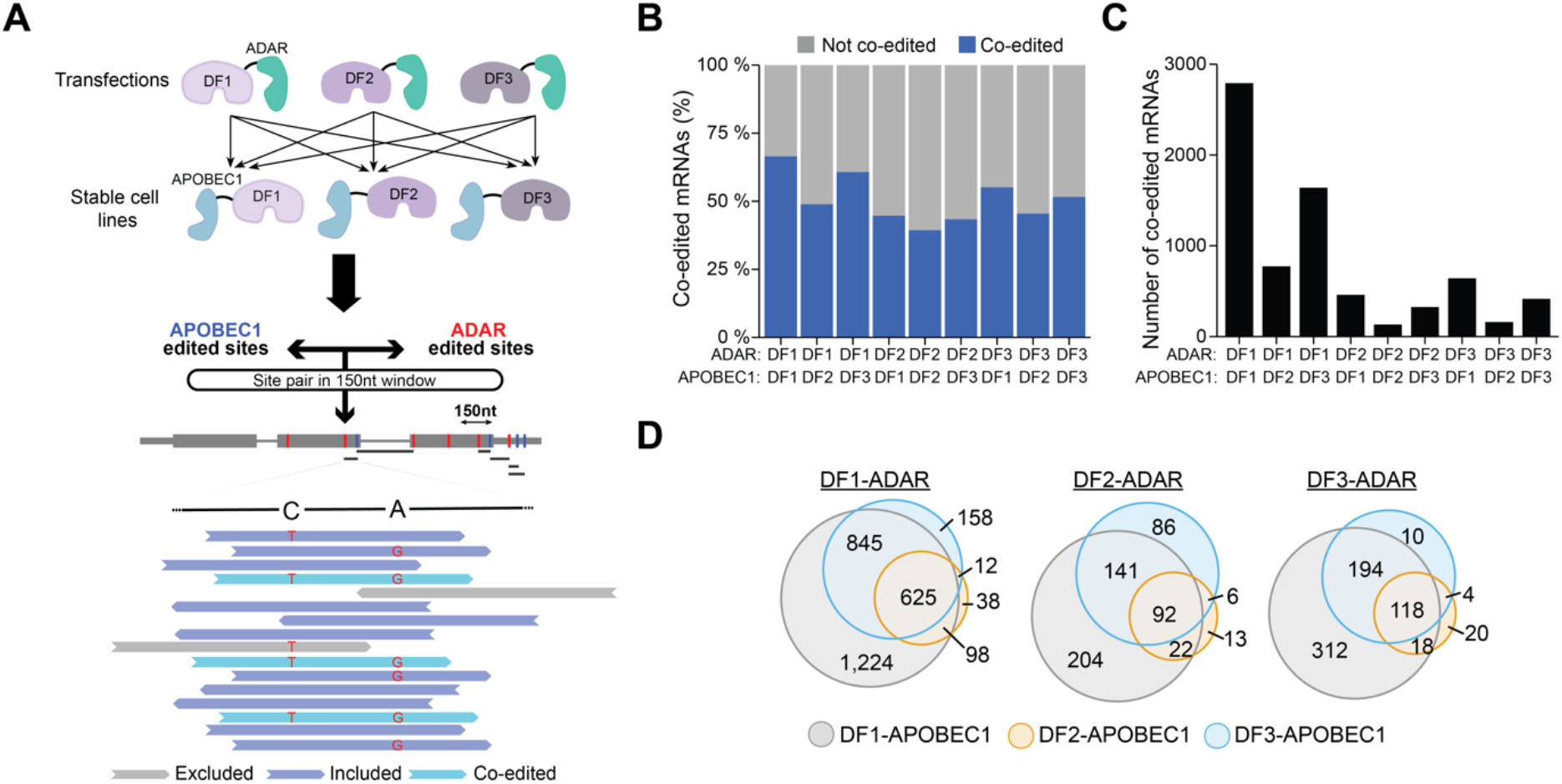
TRIBE-STAMP reveals frequent binding of individual mRNA molecules by more than one DF protein. **A**, Co-expression of DF-ADAR and DF-APOBEC1 in cells enables simultaneous detection of the target mRNAs of distinct DF proteins. Schematic shows the workflow for single-molecule TRIBE-STAMP analysis. **B**, Proportion of shared target mRNAs with single-molecule co-editing (A2I and C2U) in cells expressing the indicated DF-ADAR and DF-APOBEC1 fusion proteins. Only the shared targets with A2I and C2U editing sites within a 150 nt window on individual reads were considered. **C**, Number of mRNAs with single-molecule co-editing (A2I and C2U) in cells expressing the indicated DF-ADAR and DF-APOBEC1 fusion proteins. **D**, Target mRNAs are co-edited across different combinations of co-expressed DF fusion proteins. Euler plots show the overlap of mRNAs identified as having single-molecule co-editing when each DF-ADAR protein is co-expressed with distinct DF-APOBEC1 proteins.

We first examined TRIBE-STAMP data from cells expressing the ADAR- and APOBEC1-tethered version of the same DF protein. For all three DFs, we found that a substantial fraction of target mRNAs shared by both fusion proteins has co-editing at the single-molecule level (66.3% for DF1, 39.0% for DF2, and 51.7% for DF3) (Fig. 2B,C). The higher proportion of co-edited mRNAs in DF1-ADAR/DF1-APOBEC1-expressing cells is unlikely to be caused by the greater number of total editing sites per mRNA in these cells (Supplemental Fig. S2E), as our analysis of co-editing accounts for differences in editing frequency and sequencing coverage across datasets (see Methods). Thus, these data suggest that a large fraction of mRNAs that are recognized by a given DF protein can be bound more than once by that same DF protein on individual mRNA molecules.

We next examined cells co-expressing two different DF proteins. Surprisingly, we found that over 40% of the shared targets of any two DF proteins exhibit co-editing of the same mRNA molecules, an effect that was observed regardless of which protein was tagged with ADAR or APOBEC1 (Fig. 2B,C). The proportion of co-edited mRNA molecules was highest for cells co-expressing DF1 and DF3 (59.9% for DF1-ADAR/DF3-APOBEC1-expressing cells and 54.5% for DF3-ADAR/DF1-APOBEC1-expressing cells). We also found that many of the mRNAs co-edited at the single-molecule level are co-edited across different pairs of DF proteins, suggesting that co-binding of individual mRNA molecules is not restricted to a particular DF protein pair (Fig. 2D and Supplemental Fig. S3A). Interestingly, however, target mRNAs shared by DF2 have the lowest proportion of co-editing at the single-molecule level, indicating that DF2-bound mRNA molecules are less likely to be co-bound by DF1 or DF3 (Fig. 2B,C).

We considered the possibility that co-editing of individual RNAs could result from protein:protein interactions between two DF proteins, such that only one DF protein actually binds to the mRNA, but interacts with a second DF protein and brings it close enough to the mRNA for co-editing to occur. To explore this, we performed co-IPs in the presence and absence of RNase to determine whether the DF proteins interact with each other in an RNA-dependent or RNA-independent manner. However, we failed to observe either direct or RNA-mediated interactions between any of the DF proteins, suggesting that co-editing is due to direct binding of DF proteins to target RNAs (Supplemental Fig. S3B). Altogether, our data indicate that individual mRNA molecules can be bound by more than one DF protein throughout their lifetime. Additionally, the proportion of co-edited target mRNAs revealed by TRIBE-STAMP is likely to be an underestimate, since over 85% of possible co-edited targets for each DF protein pair have evidence for single-molecule co-editing in at least one biological replicate and since we restricted our analysis to 150 nt regions (Supplemental Fig. S3C,D).

If DF proteins largely act to degrade their target transcripts, we would expect that co-editing rates on individual reads would be either stochastic or lower than expected by chance, since the first DF binding event would trigger mRNA degradation and therefore reduce the likelihood that a second DF protein would be able to bind to that same mRNA molecule. To investigate this, we performed a quantitative analysis of co-editing frequency on individual reads for each DF protein pair. We established a rate of expected co-editing for all pairs of A2I and C2U sites within 150 nt of each other and performed a permutation test to determine whether the rate of co-editing differs significantly from the rate expected by chance (see Methods). We found that the observed co-editing rates are consistently higher than those expected by chance, an effect which was true for all combinations of DF-ADAR and DF-APOBEC1 fusion proteins (Fig. 3A and Supplemental Fig. S3E). In addition, for all DF fusion protein pairs, %A2I and %C2U editing rates were significantly higher in co-edited reads than in all reads, indicating that the presence of A2I editing increases the likelihood of also having C2U editing on the same read and vice versa (Fig. 3B). Altogether, these data indicate that binding of multiple DF proteins to the same mRNA molecule is common and occurs more frequently than expected by chance, which is consistent with a model in which DF proteins do not immediately act to degrade their target mRNAs.

**Fig. 3:**
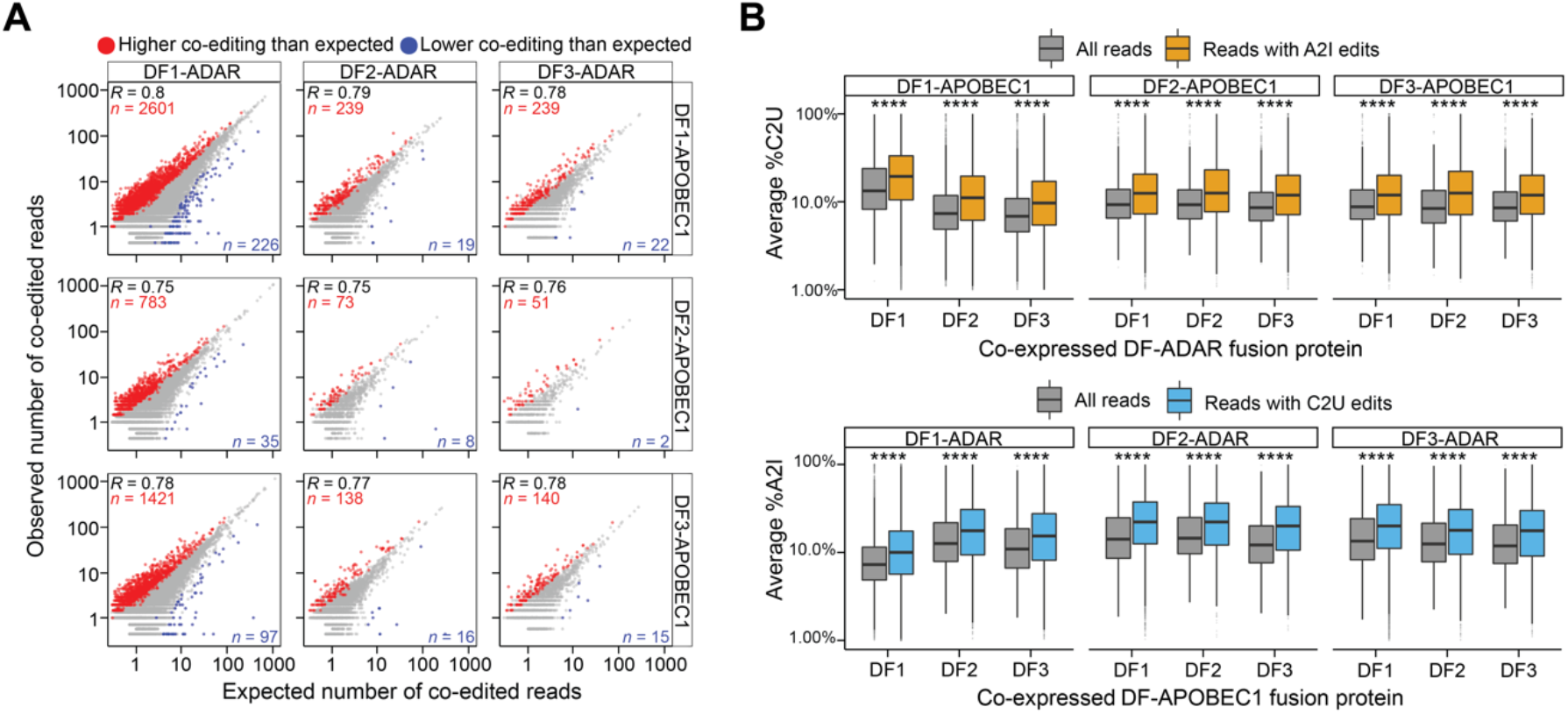
Quantitative analysis of TRIBE-STAMP co-editing. **A**, Scatterplot showing the number of expected and observed co-edited reads for all combinations of DF fusion proteins in TRIBE-STAMP datasets. Co-edited read counts that are higher or lower than expected by chance are colored in red and blue, respectively. Significance was tested using 10,000 iterations of a permutation test adjusted to 10% FDR. **B**, Boxplots showing the editing rates in TRIBE-STAMP reads that span pairs of A2I and C2U editing sites for each combination of DF fusion proteins. The editing rates in all reads (gray), in reads containing A2I edits (orange), or in reads containing C2U edits (blue) are shown. Top row: %C2U level when DF-APOBEC1 proteins are co-expressed with each DF-ADAR protein. Bottom row: %A2I when DF-ADAR proteins are co-expressed with each DF-APOBEC1 protein. Significance is the result of a Wilcoxon rank sum test adjusted for multiple comparisons. ****: *P* < 1e-5.

### RIP-TRIBE validates the findings of TRIBE-STAMP and reveals sequential binding of target mRNAs by different DF proteins

We next wondered if co-editing of the same reads reflects binding of DF proteins to two distinct m^6^A sites in close proximity on the same mRNA molecule, or whether it results from sequential binding of different DF proteins to the same m^6^A site (Supplemental Fig. S3F). To investigate this, we combined RNA immunoprecipitation (RIP) with HyperTRIBE (RIP-TRIBE) to isolate the mRNAs bound by each DF protein in cells expressing DF1, DF2, or DF3 fused to ADAR (Fig. 4A). This strategy enabled us to determine whether the target mRNAs of an individual DF protein (identified by RIP) have already been bound by a different DF protein (indicated by A2I editing). To perform RIP-TRIBE, we first generated stable cell lines expressing inducible DF1,2, or 3-ADAR-T2A-EGFP (Supplemental Fig. S4A,B). We then transfected each cell line with FLAG-tagged DF1,2 or 3 and treated cells with doxycycline for 24h to induce DF1,2, or 3-ADAR-T2A-EGFP expression. Finally, EGFP positive cells were isolated by flow cytometry and subjected to RIP using an anti-FLAG antibody. The RNAs bound by DF proteins were then identified by RNA-seq (Fig. 4A).

**Fig. 4:**
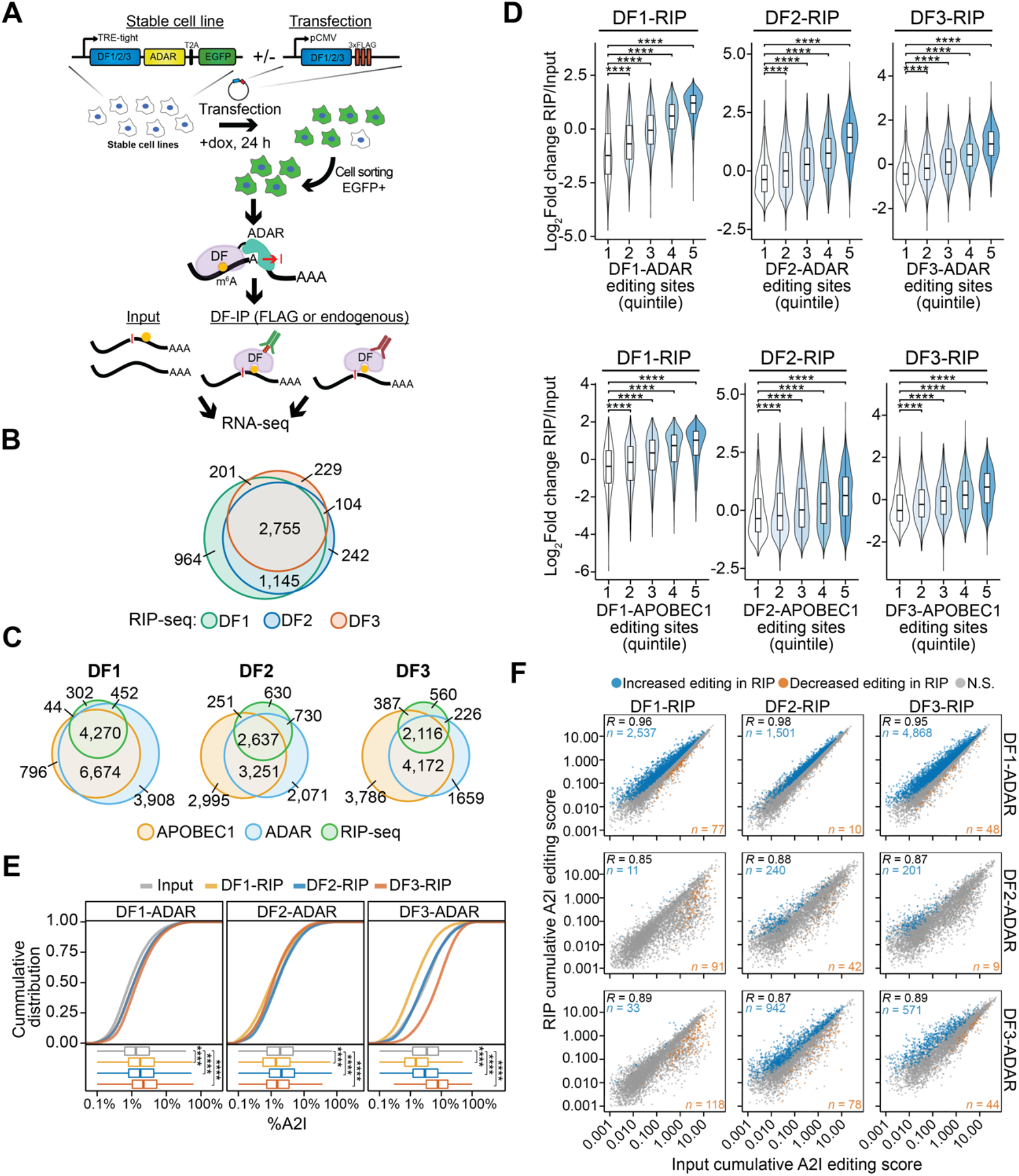
RIP-seq confirms co-binding of DF proteins to the same mRNA molecules. **A**, Overview of the RIP-TRIBE experimental workflow. Stable HEK293T cell lines expressing DF-ADAR-HA-T2A-EGFP were transfected with FLAG-tagged DF proteins, and EGFP+ cells were sorted. DF proteins were then immunoprecipitated and bound RNA was extracted and subjected to RNA-seq and A2I editing site analysis. RIP-TRIBE was performed using antibodies against either FLAG (to detect FLAG-DF protein targets) or DF proteins (to detect endogenous DF protein targets). **B**, Overlap of DF target mRNAs identified by RIP-seq. **C**, Overlap of the target mRNAs of each DF protein identified by RIP-seq and TRIBE-STAMP. DF-ADAR and DF-APOBEC1 targets are indicated. **D**, Violin plots showing the relative enrichment of mRNAs in RIP-seq datasets (RIP/Input). mRNAs are divided into quintiles based on the number of editing sites they contain in cells expressing the indicated DF-ADAR or DF-APOBEC1 proteins. Indicated *p*-values are the result of a Wilcoxon rank-sum test adjusted for multiple comparisons. ****: *P* < 1e-5. **E**, Cumulative distribution plots (top) and boxplots (bottom) showing the average A2I editing rates in mRNAs from the input and RIP fractions for each DF-RIP sample. The DF-ADAR cell line in which each RIP was performed is indicated on top. Significance was determined using a Kruskal-Wallis test followed by a post-hoc Dunn’s test adjusted for multiple comparisons. ****: *P* < 1e-5. **F**, Scatterplot showing the cumulative RNA editing scores in input and RIP samples for each DF protein. The DF protein subjected to RIP is indicated at the top, and the DF-ADAR cell line in which RIP was performed is indicated on the right. For each comparison, the mRNAs with a statistically significant increase or decrease in editing levels are colored in blue and orange, respectively. Statistical significance was determined using a Wald-test with FDR < 0.05. The Pearson correlation coefficient is indicated in the top left corner. The numbers of mRNAs with increased or decreased editing are indicated in the top left and bottom right corners, respectively.

We observed a high degree of overlap of the mRNA targets of all three DF proteins identified by RIP-seq (Fig. 4B). Additionally, the majority of mRNA targets uncovered by RIP-seq were also identified by TRIBE-STAMP (Fig. 4C), and the relative enrichment of mRNA targets in RIP samples compared to input samples correlates positively with the number of A2I and C2U editing sites in TRIBE-STAMP data (Fig. 4D). These results further validate the accuracy of the TRIBE-STAMP approach and confirm that both RIP-seq and TRIBE-STAMP can accurately identify DF protein target mRNAs.

We next compared the level of A2I editing in the input and RIP fractions for each pair of DF proteins. We reasoned that, if the target mRNAs of a given FLAG-DF protein had been previously bound by the co-expressed DF-ADAR protein, the A2I editing rate of those mRNAs would be higher in the RIP fraction than the input fraction. In contrast, if the DF-ADAR protein did not previously bind to the target mRNAs, the A2I editing rate in the RIP fraction would be lower than in the input fraction. To quantify this, we computed an editing score for each mRNA which measures the cumulative editing frequency of all called A2I sites in the transcript and which facilitates comparisons between samples by accounting for differences in coverage (see Methods). We found that transcript editing scores, as well as average %A2I values, were higher in the RIP samples compared to input for cells co-expressing FLAG- and ADAR-tagged versions of the same DF protein (Fig. 4E,F). This is consistent with our TRIBE-STAMP data indicating that mRNAs can be bound more than once by the same DF protein (Fig. 2B,C).

We next examined A2I editing in RIP samples from cells expressing FLAG- and ADAR-tagged versions of different DF proteins. We found that average %A2I in the RIP fractions of all three DF proteins was higher than the input fraction in DF1-ADAR-expressing cells (Fig. 4E). In contrast, average %A2I in DF2-ADAR- and DF3-ADAR-expressing cells was only higher than input in the DF2 or DF3 RIP samples, respectively (Fig. 4E). Together, these data suggest that the mRNAs bound by DF2 or DF3 are likely to have been previously bound by DF1. Indeed, we identified thousands of mRNAs that have significantly higher editing scores in DF1, DF2, or DF3 RIP samples compared to input in DF1-ADAR-expressing cells (Fig. 4F; Supplementary Table S4). Furthermore, we found that editing scores of mRNAs edited by DF3-ADAR are enriched in the RIP fraction of DF2 but not DF1 (Fig. 4F). This indicates that many DF2-bound transcripts have been previously bound by DF3, whereas DF1-bound transcripts have not. Additionally, we found that few transcripts have enriched editing in the DF1 and DF3 RIP fractions from DF2-ADAR-expressing cells, suggesting that most mRNAs bound by DF1 or DF3 have not been previously bound by DF2 (Fig. 4F). This is consistent with our TRIBE-STAMP data showing the lowest levels of single-molecule co-editing for DF2 targets (Fig. 2B,C) and may reflect a greater propensity for DF2-bound mRNAs to be targeted for degradation.

To validate these observations, we repeated the RIP-TRIBE experiments using antibodies specific to each endogenous DF protein. As before, we confirmed the overlap of target mRNAs among the three DF proteins (Supplemental Fig. S4C), and we found a positive correlation between the number of TRIBE and STAMP editing sites and the relative enrichment of mRNA in RIP samples (Supplemental Fig. S4D). Importantly, endogenous DF RIPs validated our finding that DF1-ADAR edited mRNAs were more frequently edited in DF1,2 or 3 RIP samples (Supplemental Fig. S4E; Supplementary Table S5). Additionally, we performed co-IPs detecting both endogenous and ADAR-tagged DF proteins in each DF-ADAR cell line and found no evidence of direct interactions between any of the DF proteins, which is in agreement with what we observed in wild type cells and indicates that the increased editing in RIP fractions is not due to indirect editing events (Supplemental Fig. S4F,G). Overall, our RIP-TRIBE data from endogenous DF proteins validates what we observed using FLAG-tagged DFs and supports a model in which DF proteins bind sequentially to target mRNAs.

Since DF proteins bind to m^6^A, one prediction of this model is that regions of co-editing in TRIBE-STAMP data should contain only one m^6^A site. Indeed, when we examined extended regions surrounding co-edited sites, we found that the vast majority contain only one m^6^A site (Supplemental Fig. S5A). These regions are also no more likely than non-co-edited regions to contain the m^6^A consensus motif (DRACH), suggesting that they are not inherently prone to hypermethylation (Supplemental Fig. S5B). Furthermore, the stoichiometry of most m^6^A sites is generally low (Liu et al. 2013; Molinie et al. 2016; Garcia-Campos et al. 2019; Tegowski et al. 2022), and m^6^A profiling in single cells shows that the majority of m^6^A sites do not occur within the same cell (Tegowski et al. 2022). Thus, there is a low likelihood of any two individual m^6^A residues occurring in proximity on the same mRNA molecule. Collectively, these data suggest that co-editing of individual mRNA molecules reflects sequential binding of different DF proteins to the same m^6^A site as opposed to binding to distinct m^6^A sites that occur close to each other. mRNAs edited by DF1 are those with the highest likelihood of being bound by another DF protein, whereas mRNAs edited by DF2 or DF3 are less likely to be bound by DF1. These observations are consistent with a model in which DF1 binds to methylated mRNAs first, followed by DF2 or DF3. Moreover, the finding that DF2-bound transcripts are the least likely to be subsequently bound by DF1 or DF3 suggests that DF2 targets may be more likely to undergo rapid degradation after DF2 binding.

### Polysome fractionation coupled with HyperTRIBE reveals that DF-bound mRNAs are not preferentially translated

Our finding that DF1 and DF3 bind to mRNAs before DF2 suggests that these proteins may have functions other than promoting mRNA decay. Previous studies in neurons and HeLa cells have shown that DF1 and DF3 promote mRNA translation (Wang et al. 2015; Li et al. 2017; Shi et al. 2017; Shi et al. 2018; Weng et al. 2018), although other studies in HeLa cells and mESCs have suggested that DF proteins have no direct impact on translation (Lasman et al. 2020; Zaccara and Jaffrey 2020). To gain insights into this, we reasoned that we could couple polysome fractionation with analysis of A2I editing scores in cells expressing each DF-ADAR protein to determine whether mRNAs bound by DF proteins are differentially translated compared to non-target mRNAs.

We subjected stable cells expressing DF1-ADAR, DF2-ADAR, or DF3-ADAR to polysome fractionation followed by RNA-seq (Fig. 5A; Supplemental Fig. S6A-E). We then compared A2I editing rates of mRNAs in the input and polysome fractions for each cell line to determine whether transcripts bound by individual DF proteins are preferentially localized to polysomes. We found that nearly all target mRNAs of each of the three DF proteins can localize to polysomes (DF1: 99.0%, DF2: 99.2%, DF3: 96.8%) (Fig. 5B). However, only approximately 60% of the targets of each DF protein that are present in the polysome fraction are edited there, and we found no preferential enrichment of polysome-edited targets for any of the DF proteins (DF1: 61.0%, DF2: 62.2%, DF3: 61.9%) (Fig. 5C). In addition, the average A2I editing rates of target mRNAs are slightly lower in polysome fractions compared to the input fractions for all three DF proteins, indicating that binding by DF proteins does not cause preferential enrichment of target mRNAs to polysomes (Fig. 5D). Indeed, only a small number of transcripts has increased or decreased editing in polysome fractions relative to input (Fig. 5E and Supplemental Table S6). Interestingly, although the majority of endogenous and ADAR-fused DF1, DF2, and DF3 proteins are localized to the free RNP fraction, a substantial amount of DF1 and DF2 is associated with polysomes, whereas DF3 is largely excluded (Fig. 5A and Supplemental Fig. S6A-C). Taken together, these data indicate that the majority of DF1, DF2, and DF3 target mRNAs are localized at some point in their lifetime to polysomes, but that binding by DF proteins does not commit these mRNAs to preferential polysome localization. These findings are consistent with a limited role for DF proteins in promoting mRNA translation in HEK293T cells and suggest that no single DF protein preferentially drives translation of its target mRNAs.

**Fig. 5:**
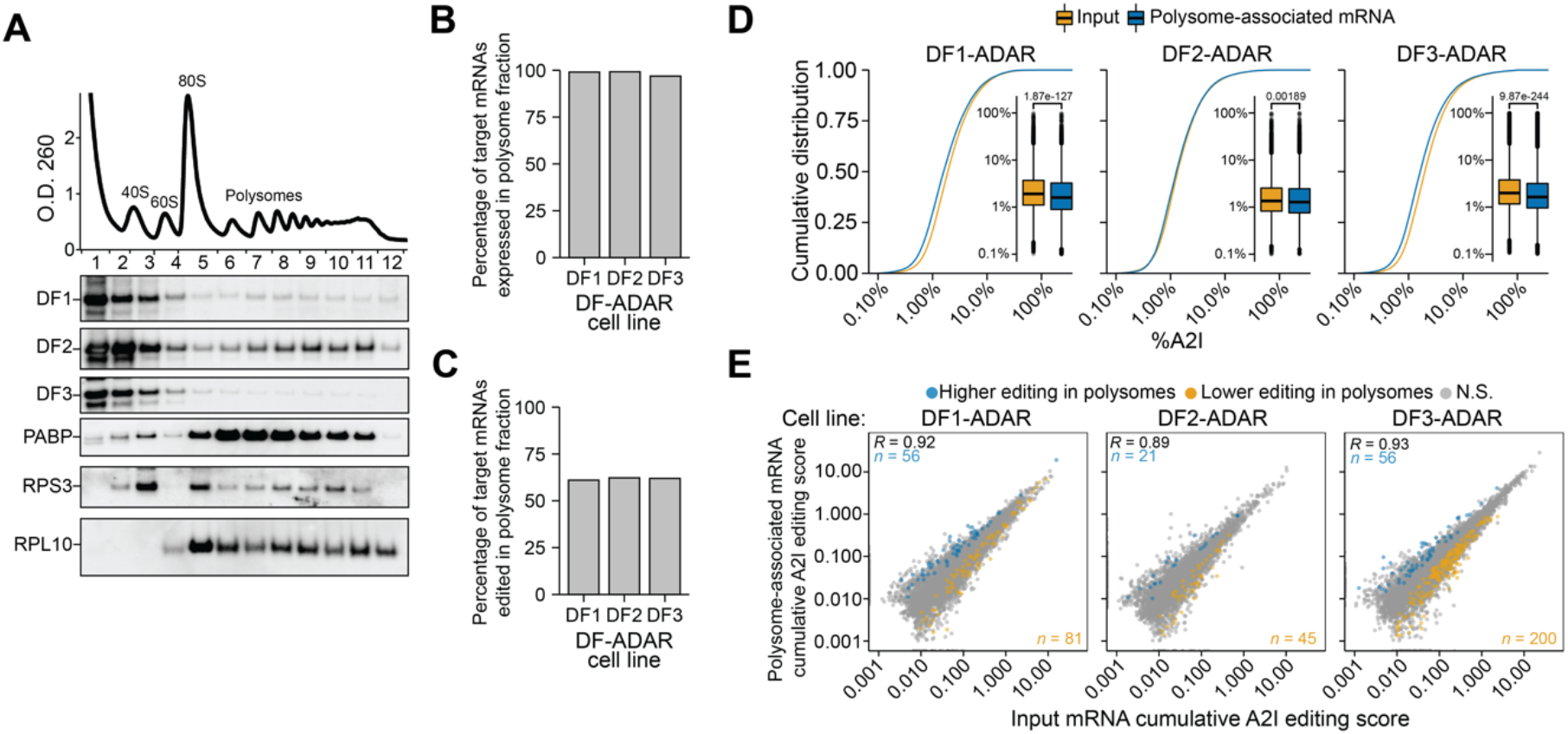
Binding by DF proteins does not preferentially partition mRNAs to polysomes. **A**, DF1 and DF2 are associated with polysomes. Representative O.D.260 signal trace during polysome fractionation of cells expressing DF-ADAR proteins is shown at the top. Western blot for individual endogenous DF proteins is shown below. Western blots for PABP and ribosomal proteins RPS3 and RPL10 are also shown as validation of polysome fractionation. **B**, Proportion of the target mRNAs of each DF protein that are detected (FPKM ≥1) in polysome-associated RNA fractions. **C**, Proportion of the target mRNAs of each DF protein that are edited in the polysome-associated fraction. **D**, Cumulative distribution plots and boxplots of the A2I editing levels in input and polysome-associated mRNAs for each DF-ADAR cell line. *P*-value is the result of a 2-sided Wilcoxon rank sum-test. **E**, Scatterplot of the cumulative RNA editing scores in input and polysome samples for each DF-ADAR cell line. mRNAs with statistically significant increased or decreased editing levels in polysome-associated RNA fractions are colored in blue and orange, respectively. The statistical significance was determined using a Wald-test, FDR < 0.05. The Pearson correlation coefficient is indicated in the top left corner. The numbers of mRNAs with increased or decreased editing are indicated in the top left and bottom right corners, respectively.

## Discussion

We present here an experimental and computational workflow called TRIBE-STAMP, a method for simultaneous detection of the RNA targets of distinct RBPs in cells. By leveraging the ability of ADAR- and APOBEC1-tethered RBPs to induce distinct mutation signatures in target RNAs, TRIBE-STAMP offers a simple, robust method for identifying the target RNAs of two RBPs at once in cells and for dissecting RNA:protein interactions at the molecular level. This approach therefore overcomes some of the major limitations of RIP- and CLIP-based methods, which focus on a single RBP at a time and which do not enable analysis of RBP binding to individual RNA molecules.

To demonstrate the utility of the TRIBE-STAMP method, we used it to address an area of recent discrepancy in the epitranscriptomics field, which pertains to the function and targets of the YTHDF m^6^A reader proteins. Two opposing models have emerged for the function of these proteins: one in which they can bind unique subsets of cellular mRNAs and carry out distinct functions, and another in which they bind to the same mRNAs and function redundantly to promote mRNA degradation. One factor contributing to the discrepancy between these two models is whether DF proteins bind different mRNAs or the same mRNAs. To address this question at the single-molecule level, we applied TRIBE-STAMP to each combination of DF protein pairs. This enabled identification of the mRNA targets of distinct DF proteins in the same cells and allowed us to examine for the first time whether individual mRNA molecules can be targeted by more than one DF protein.

TRIBE-STAMP revealed that the majority of target mRNAs of each DF protein are also bound by the co-expressed DF protein, regardless of the DF protein pair being examined, a finding that we validated by RIP-seq against both tagged and endogenous DF proteins. However, we also identified many mRNA targets that are unique to DF1 compared to DF2 or DF3, which mirrors what we found when we reanalyzed DF iCLIP data from HeLa cells (Patil et al. 2016), as well as our previous RIP-seq studies in hippocampal neurons (Flamand and Meyer 2022). Overall, our data demonstrate that TRIBE-STAMP can accurately identify the target RNAs of two RBPs simultaneously in cells and showed that the mRNA targets of all three DF proteins are largely shared at the gene level.

Binding of all three DF proteins to the same target mRNAs has been used to support the functional redundancy model, in which all DF proteins promote the degradation of their target mRNAs (Kontur et al. 2020; Lasman et al. 2020; Zaccara and Jaffrey 2020). One prediction of this model is that individual mRNA molecules are bound by only one DF protein during their lifetime, since they are then targeted for degradation. Surprisingly, our single-molecule TRIBE-STAMP analysis revealed that individual copies of a given mRNA are bound by multiple DF proteins throughout their lifetime. We also found that individual mRNA molecules can be bound by the same DF protein more than once in their lifetime. Although these findings do not rule out a role for each of the DF proteins in promoting mRNA degradation, they suggest that DF proteins do not promote degradation every time they bind to a target mRNA (Fig. 6). This could be caused by inefficient recruitment of the degradation machinery, or it could reflect transient binding of DF proteins to their target mRNAs, such that rapid dissociation from target transcripts does not always allow sufficient time to elicit degradation. Indeed, binding affinities of the YTH domain to m^6^A are in the high nanomolar range (Wang et al. 2014; Xu et al. 2015; Liu et al. 2018), which is on the low end of RNA:protein interaction affinities (Yang et al. 2013; Harini et al. 2022). In this sense, the idea that all three DF proteins promote degradation is fitting, since it may be a failsafe in case one DF protein binds and dissociates before it can promote degradation. Future studies examining the dwell time of individual DF proteins on their target mRNAs would likely provide more insight.

**Fig. 6:**
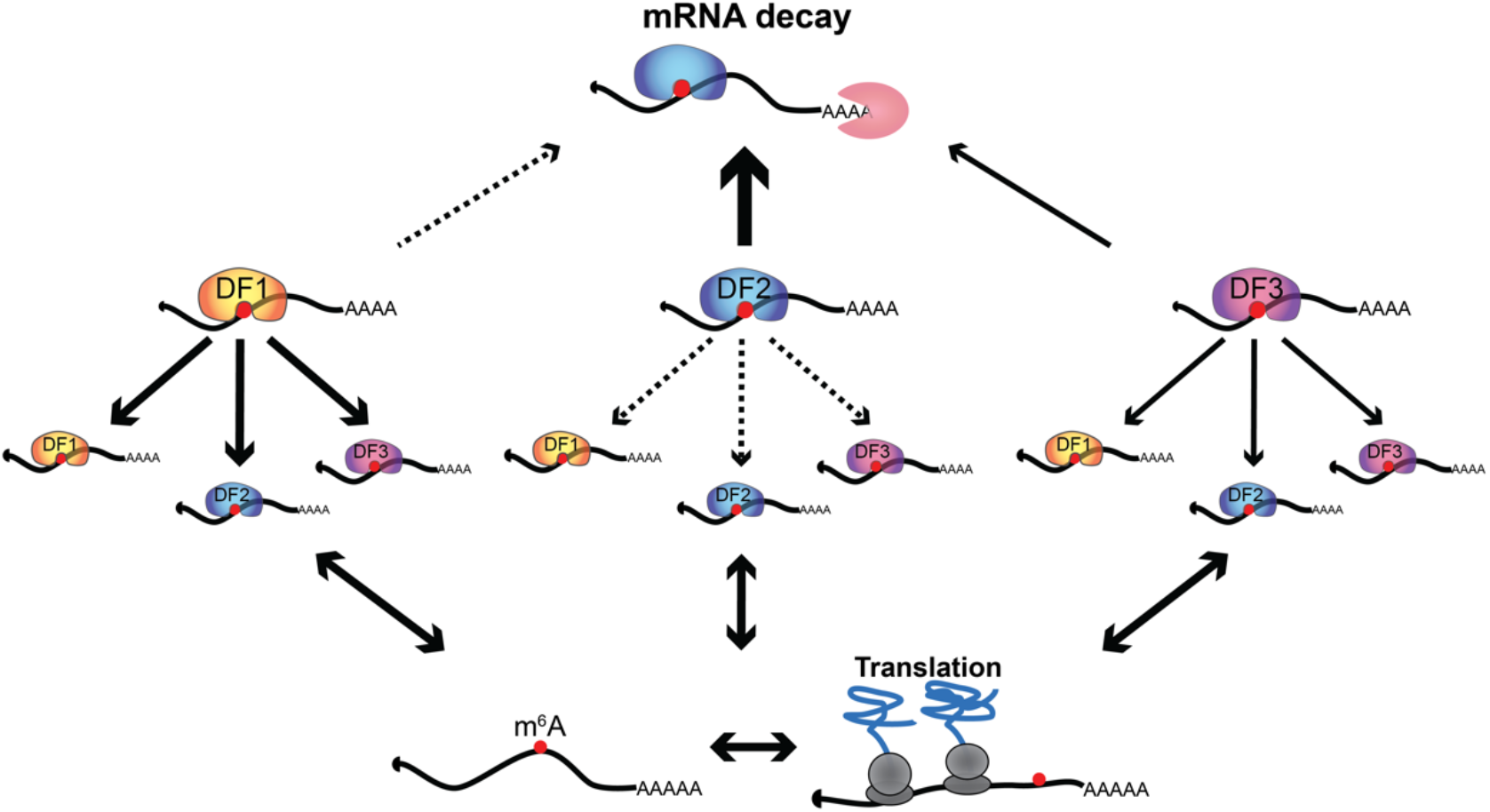
Model for the recognition of m^6^A by DF proteins. Most m^6^A-methylated mRNAs are shared targets of DF1, DF2 and DF3 proteins, and individual mRNA molecules can be bound by more than one DF protein in their lifetime. DF1 generally binds before DF2 or DF3 and does not lead to rapid mRNA decay. In HEK293T cells, binding by DF proteins does not lead to preferential recruitment of mRNAs to polysomes.

The majority of mRNAs in our TRIBE-STAMP data contain only one m^6^A site in the vicinity of co-edited regions, indicating that the co-editing of individual mRNA molecules that we see for pairs of DF proteins is unlikely to reflect binding to more than one m^6^A site. This is also supported by previous studies which have shown that most m^6^A sites are present at low stoichiometry and that clusters of m^6^A sites occur rarely within the same cell (Liu et al. 2013; Molinie et al. 2016; Garcia-Campos et al. 2019; Tegowski et al. 2022). Thus, the likelihood that more than one m^6^A site is found on the same mRNA molecule is very low. Instead, we find that co-editing likely reflects sequential binding by distinct DF proteins to the same m^6^A sites. Using RIP-TRIBE, we demonstrate that the target mRNAs immunoprecipitated by DF1, DF2, and DF3 are enriched for A-to-I editing when the RIPs are performed in cells expressing DF1-ADAR, indicating that these mRNAs were previously bound by DF1. This suggests a model in which DF1 binds to target mRNAs before DF2 and DF3. We observed a similar effect for mRNAs immunoprecipitated by DF2 in DF3-ADAR cells, indicating that DF2-bound transcripts can be previously bound by DF3. These results align with previous studies that used 4SU labeling of nascent RNA coupled with DF RIP and found that DF1 and DF3 bind to RNAs before DF2 (Shi et al. 2017). One possibility is that DF2 more efficiently promotes mRNA degradation compared to DF1 and DF3, therefore making it more likely that DF2-bound transcripts are less available for subsequent binding by DF1 or DF3 (Fig. 6). This would be consistent with our finding that the mRNAs actively bound by DF1 and DF3 (as indicated by RIP) are less likely to have been previously bound by DF2 (as indicated by DF2-ADAR-mediated A-to-I editing).

Our findings that DF1 binds more target mRNAs than DF2 or DF3 and that DF1 binds mRNAs before DF2 and DF3 led us to hypothesize that DF1 may have functions other than mRNA degradation. DF1 has previously been shown to promote mRNA translation (Wang et al. 2015; Shi et al. 2018; Zhuang et al. 2019), so we explored whether DF1-bound transcripts are preferentially localized to polysomes. We were surprised to find that mRNAs edited by DF1-ADAR show similar levels of editing enrichment in polysomes as those edited by DF2- or DF3-ADAR. Although we do not know whether editing of these polysome-associated mRNAs reflects active or previous binding by DF-ADAR proteins, these data suggest that DF1 is not preferentially promoting the localization of its target transcripts to polysomes compared to DF2 or DF3 in HEK293T cells. It is notable that DF1 has been shown to promote translation in other cell types and unique cellular contexts, including in neurons in response to activity and axonal injury (Shi et al. 2018; Weng et al. 2018; Zhuang et al. 2019), so it may be that the translation-promoting effects of DF1 are cell type- and context-specific.

Here, we used TRIBE-STAMP to assess the binding of YTHDF proteins to cellular mRNAs, but we expect that this method can be applied to any RBP pair of interest to identify shared transcripts and single-molecule binding within the same cells. Additionally, we used stable cell lines expressing ADAR- and APOBEC1-tagged proteins at similar levels to their endogenous counterparts, but tagging of endogenous RBP loci with ADAR and APOBEC1 is an alternative approach that could be used. Furthermore, TRIBE-STAMP could be coupled with long-read sequencing to identify RNA molecules co-bound by RBPs that recognize distinct transcript regions separated by large distances, such as the 5’UTR and 3’UTR. In theory, TRIBE-STAMP could also be used in conjunction with PUP-2 or APEX2 labeling to identify the targets of more than two RBPs at once (Lapointe et al. 2015; Fazal et al. 2019; Padron et al. 2019; Xu et al. 2022). Thus, the TRIBE-STAMP method provides a versatile approach for investigating RNA:protein interactions in cells and enables single-molecule binding information for multiple RBPs at once, therefore overcoming the limitations of many current protein-centric methods which only examine single RBPs in isolation.

## Acknowledgments

We thank members of the Meyer laboratory for scientific discussion and helpful commentary. We are grateful to the Duke Center for Genomic and Computational Biology, the Duke Molecular Genomics Core, and the Duke Human Vaccine Institute Flow Cytometry Core for providing the infrastructure and support for sequencing and cell sorting. This research was supported by the Canadian Institutes of Health Research (M.N.F.) and the National Institutes of Health (R01MH118366 and DP1DA046584 to K.D.M.). K.D.M. is also supported by the Rita Allen Foundation.

## Author Contributions

K.K., M.N.F., and K.D.M. designed experiments. M.N.F. designed the analysis pipeline and performed the bioinformatic data analysis described in the manuscript. K.K. and M.N.F. carried out the experiments with additional contributions from R.T.. M.N.F., K.K., and K.D.M. interpreted data and wrote the manuscript.

## Methods

### Employed cell lines

HEK293T cells (ATCC, Manassas, VA) were maintained at 37 °C and 5% CO_2_, using Dulbecco’s Modified Eagle’s Medium (Corning, Corning, NY) supplemented with 10% fetal bovine serum (Avantor, Radnor, PA) and 10 units/mL Penicillin, and 10 μg/mL Streptomycin (Gibco, Waltham, MA).

### Generation of stable cell lines

HEK293T cells at ~50% confluency were transduced with filtered lentiviral-containing supernatant for 24 h. Stable cell lines were selected by adding 2 μg/mL puromycin in fresh growth medium for 7 days before further analysis. All HEK293T transgenic cell lines were plated 2 days before the experiment. The following day, the cells were treated with 1 μg/mL doxycycline (Sigma-Aldrich, St. Louis, MO).

#### Antibodies

The following antibodies and concentrations were used: rabbit anti-GAPDH (Proteintech; 10494-1-AP; WB-1:1,000); rabbit anti-HA (CST; C29F4; WB-1:1,000); rabbit anti-ADARB1 (Aviva Systems; OAAF02345; WB-1:1,000); mouse anti-FLAG M2 (Sigma, F1804, WB-1:2,000, IP: ?); rabbit anti-YTHDF1 (Abcam; ab252346 ;WB-1:1,000); rabbit anti-YTHDF2 (Abcam; ab246514 ;WB-1:2,000), rabbit anti-YTHDF3 (Abcam; ab220161 ;WB-1:2,000); Rabbit anti-RPL10 (GeneTex, GTX55169, 1:1000); Rabbit anti-RPS3 (CST, 2579S, 1:1000); Rabbit anti-PABP (CST, 4992S, 1:2000); horseradish peroxidase (HRP)-conjugated goat anti-rabbit (Abcam; ab6721; 1:2,500); HRP-conjugated goat anti-mouse (Invitrogen; 62-6520; 1:10,000).

### Western blotting

Protein samples were separated on 4–12% NuPAGE gels (Thermo), at 175V for 60 min and transferred to nitrocellulose membranes at 100 V for 90 min in transfer buffer (25 mM Tris-Cl pH 8.3, 192 mM glycine, 20% methanol). Blocking was carried out for ≥30 min in 5% nonfat dry milk in 0.1% PBST and antibodies were incubated overnight in blocking buffer or 5% BSA at 4°C. Secondary antibodies were incubated on membranes in blocking buffer for 1 h at room temperature. ECL reagent (Amersham ECL Prime) was mixed 1:1 and added to the membranes, which were imaged using the Bio-Rad Chemidoc MP imaging system.

### DF protein expression and RNA isolation for TRIBE-STAMP

Expression of APOBEC1-DF-HA-T2A-EGFP in HEK293T stable cells was induced by 1 μg/mL of doxycycline for 24 h. Plasmids pcDNA3-HA-DF-hADARcd-E488Q-T2A-mCherry were transfected into HEK293T stable cell lines of APOBEC1-DF-HA-T2A-EGFP using Fugene HD. 24 or 48 h after transfection, more than 500,000 EGFP-mCherry-double-positive cells were sorted and collected using a BD FACSAria II. Total RNA was extracted from the sorted cells with TRIzol LS reagent, followed by DNase I treatment, phenol:chloroform extraction and ethanol precipitation. RNA quantity and quality were measured using the Qubit RNA high sensitivity assay and the RNA 6000 Pico assay on the Bioanalyzer 2100. Samples with a RIN >= 7 were used for library preparation.

### RNA-seq library construction and sequencing

RNA-seq libraries for TRIBE-STAMP were prepared from 0.5 μg of total RNA in triplicates (DF1-ADAR and DF2-ADAR) or duplicates (DF3-ADAR) for each co-expressed DF-APOBEC1 protein using the NEBNext Ultra II Directional RNA Library Prep Kit for Illumina (NEB #E7760L) with NEBNext Poly(A) mRNA Magnetic Isolation Module (NEB #E7490L). cDNA libraries were sequenced by the Sequencing and Genomic Technologies Shared Resource at Duke University on an Illumina NovaSeq6000 using a S-prime or S1 flow cell with paired-end 250- or 150-bp reads respectively, yielding 42-132M million clusters per library. Libraries for cells expressing ADAR or APOBEC1 alone were sequenced on an Illumina NovaSeq6000 using a S-prime flow cell with a 250-bp paired-end protocol.

RNA-seq libraries for RIP-TRIBE and polysome were prepared in triplicates from 150 ng of total RNA using the NEBNext Ultra II Directional RNA Library Prep Kit for Illumina with the NEBnext rRNA depletion Kit (NEB #E6310L). cDNA libraries were sequenced on an Illumina NovaSeq6000 using S-prime or S1 flow cells with paired-end 150-bp reads, yielding an average of 32.5 million clusters for FLAG-DF RIP-seq, 23.6 million clusters for endogenous DF RIP-seq and 45.8 million clusters for polysome associated RNA-seq.

### RNA immunoprecipitation

RIP-TRIBE experiments were conducted following induction of DF-ADAR-HA-T2A-EGFP expression in HEK293T stable cells with 1 μg/mL of doxycycline for 24 h. For FLAG-DF RIP experiments, pCMV-3xFLAG-DF plasmids were transfected into HEK293T stable cell lines using Fugene HD. 24 h after induction and transfection, 10 million EGFP-positive cells were sorted and collected using a BD FACSAria II. Each protein was immunoprecipitated using 4 μg of anti-FLAG M2 or anti-YTHDF1, 2, or 3 antibodies for FLAG-DF IP or endogenous DF IP, respectively. Antibodies were pre-coupled to 50 μL Dynabeads Protein A (Invitrogen, 10001D) in PBS containing 0.02% Tween and incubated at 21°C for 30 min. 10 million flow cytometry sorted cells were lysed in RIP buffer (25 mM Tris-Cl pH 7.4, 150 mM NaCl, 2.5 mM EDTA, 0.5 mM DTT, 0.5% NP-40, RNase inhibitor, Roche cOmplete mini EDTA-free protease inhibitor) by mixing for 15 min at 4°C. Lysate was cleared by centrifugation at 17,000 x g for 10 min and the supernatant was transferred to a new tube and incubated with antibody-coupled beads for 1 h at 4°C. Following immunoprecipitation, beads were washed 3 times for 5 min with washing buffer (25 mM Tris-Cl pH 7.4, 150 mM NaCl, 2.5 mM EDTA, 0.5 mM DTT, 0.1% NP-40). One-fifth of the beads were used for protein elution in 2x NuPAGE buffer with 100mM DTT, followed by western blot analysis. The rest of the beads were resuspended in 1mL of Trizol (Invitrogen) and RNA was extracted. Precipitated RNA was washed twice with 75% ethanol and resuspended in water. A fraction of the cell lysate was used for RNA input extraction and treated with TurboDNase (Invitrogen) for 30 min at 37°C.

### Polysome profiling

Linear sucrose gradients were prepared from 10% and 50% sucrose solutions (10 mM HEPES pH.7.4, 150 mM KCl, 1 mM MgCl_2_, 1 mM DTT, 100 μg/mL cycloheximide) using a Gradient Master model 108 (Biocomp) in 25 x 89 mm open top polyclear centrifuge tubes (#7052, SETON) following the manufacturer’s instructions. For each experiment, 4 x 10 cm-dish of cells were treated with 100 μg/mL cycloheximide for 10 min at 37°C and washed once in PBS containing 100 μg/mL cycloheximide. Cells were removed from the plates using a cell scrapper, pelleted at 300g for 5 min at 4°C and resuspended in 500 μL of lysis buffer (10mM HEPES pH.7.4, 150 mM KCl, 1 mM MgCl_2_, 2% NP-40, 1 mM DTT, 100 μg/mL cycloheximide, 40U/mL RNaseOUT, cOmplete Mini EDTA-free protease inhibitor (Roche)). After 10 min lysis on ice, the lysate was cleared by centrifugation at 17,500g for 10 min at 4°C and carefully overlaid on the prepared gradients. Gradients were ultracentrifuged at 28.000g for 3 h at 4°C in a SW 28 Ti rotor (Beckman). 1.13 mL fractions were then collected using the Gradient Station model 153, piston gradient fractionator model 152 (Biocomp) and a Gilson FC203B fraction collector. Absorbance at 260 nm was measured using a TRIAX flow cell (Biocomp). For RNA-seq, fractions mapping to polysomes were pooled and precipitated with 0.1 volume of 3 M sodium acetate pH 5.2, 1 μL Glycoblue co-precipitant (Thermo) and 2 volumes of 100% ethanol for 16 h at −20°C. RNA was pelleted by centrifugation at 10,000g for 30 min at 4°C, washed twice in 75% and resuspended in nuclease-free water. For western blot analysis, proteins were precipitated with 0.03% sodium deoxycholate and 10% TCA for 30 min on ice, pelleted by centrifugation at 17.500g for 10 min and washed twice with ice-cold acetone. Protein pellets were resuspended in 1x NuPAGE loading buffer with 100mM DTT in volumes adjusted by the absorbance at O.D.260 in each fraction. The RNP, 40S, 60S and 80S fractions were pooled for analysis.

### TRIBE-STAMP analysis

#### Identification of editing sites

To identify editing sites, we used Bullseye (https://github.com/mflamand/Bullseye), a set of Perl scripts adapted from the HyperTRIBE pipeline (McMahon et al. 2016), which we developed for analysis of DART-seq (Meyer 2019; Tegowski et al. 2022). BAM files were parsed to generate coverage matrices at each position in the genome, excluding duplicate and multi-mapped reads. Editing sites for STAMP or HyperTRIBE in individual samples were then called by comparing C2U or A2I mutation rates at all positions within annotated exons (hg38 GENCODE V36) to samples expressing APOBEC1 or ADAR alone. Sites edited at rates between 2.5% and 99%, edited to higher levels than control cells (1.5x for C2U; 3x for A2I), with at least 3 mutations, and which did not overlap annotated SNP (dbSNP153) were kept for further analysis. The final list of sites was identified as those found in at multiple replicates (C2U: 2/9; DF1-ADAR and DF2-ADAR: 4/9; DF3-ADAR: 3/6) and with an average editing rate of at least 5% in all samples where sites were called. To identify editing sites found in methylated regions, we merge a MeRIP-seq dataset in HEK293T cells (GSE29714) to a single nucleotide resolution miCLIP datasets in HEK293 cells (GSE63753) using bedtools. We then broadened each peak by 25-nt on each direction to identify regions of the transcriptome that likely contain m^6^A sites. Editing sites overlapping these regions was then obtained using bedtools intersect.

#### Identification of co-editing events

To quantify co-editing events, we used co_editing.pl from the Bullseye pipeline using the following options: -minCov 20 -PermutationTest 10000 -removeDup -removeMultiMapped. Briefly, this pipeline first identifies pairs of editing sites found within 150 nt of each other, allowing pairs of sites to span over one annotated intron (hg38 GENCODE V36). BAM files are then parsed to identify pairs of sites for which at least 20 reads cover both positions. For each pair, we calculated the expected co-editing rate as:

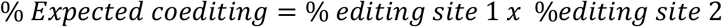

To account for differences in coverage and editing rates at individual sites, we filtered pairs of sites to only keep those with at least 3 mutations at each individual site and for which the expected co-editing frequency multiplied by the coverage is greater than one; or with at least one co-edited read:

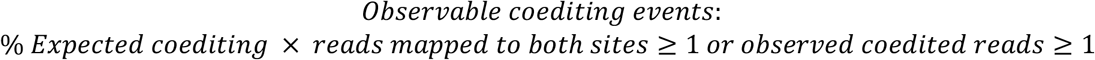

We considered a pair of sites to show exhibit co-editing when at least 2 reads were edited at both positions in all biological replicates (3 biological replicates for DF1-ADAR and DF2-ADAR-expressing cells, 2 biological replicates for DF3-ADAR-expressing cells). To identify statistically significant co-editing events, we measured the distribution of non-edited, individually edited, and co-edited reads using a chi-squared test. We then measured the probability that the observed values are more extreme than chance by comparing the observed chi-squared statistic to those obtained when the read distribution was shuffled 10,000 times based on the base editing frequency at individual sites.

#### Quantification of editing in RIP-TRIBE and polysome-associated mRNAs

To measure changes in editing across conditions, we first generated matrices containing the coverage and number of mutations at each called site for all samples of a RIP-TRIBE or polysome profiling-RNA-seq dataset. To identify sites with statistically significant changes in editing, we fitted the editing frequency at each site in a quasibinomial general linear model in R (4.1.2) using the glm function: glm(formula= mut/cov ~ condition,data=df,weights = cov, family=”quasibinomial”). In this model, we only considered samples where at least 20 reads were mapped to each editing site. The statistical significance was then measured using a Wald’s test on the fitted data and adjusted for multiple comparisons using independent hypothesis weighing (IHW) (Ignatiadis et al. 2016). To measure changes in editing at the gene level, we summed mutations and coverages of all called sites within each annotated gene and fitted the aggregate editing frequency in the quasibinomial generalized linear model. To compare editing at the RNA level, we established an RNA editing score which is defined as:

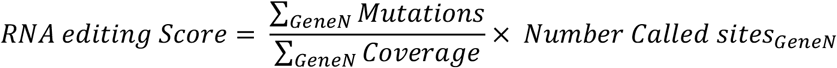

We defined mRNAs with significant changes in editing as those with at least a 1.25-fold change in editing and an adjusted p-value of less than 0.05.

#### Analysis of publicly available iCLIP datasets

Data from DF1, DF2 and DF3 iCLIP profiling in HEK293T cells (Patil et al. 2016) (GSE78030) were downloaded and used for the identification of targets of each DF protein. Raw reads were trimmed with Flexbar with the following parameters: --zip-output GZ --max-uncalled 2 --min-read-length 15 -a adapter_iCLIP.fasta -q TAIL -qf i1.8 with the AGATCGGAAGAGCGGTTCAG adapter sequence. UMIs were extracted from the trimmed reads using UMI_tools (v1.1.1) (Smith et al. 2017) by running: umi_tools extract -I file.fastq.gz --bc-pattern=NNNNNNNNN. Processed FASTQ files were then aligned to the hg38 genome using NovoAlign (v4.03.03) with the following options: -t 85 -l 16 -s 1 -o SAM -r None -a. Duplicate reads were then removed from the BAM files with UMI-tools dedup. For each DF iCLIP dataset, BAM files were parsed and peaks called using the CTK pipeline (Shah et al. 2017). Statistically significant peaks (p < 0.05) were used to identify transcripts targeted by each DF protein.

#### Statistics and reproducibility

Statistical analyses were performed using R 4.1.0. Quantitative data are represented as the mean ± s.d. or as boxplots where the center line represents the median, the box limits the 25^th^ and 75^th^ percentiles, and the whiskers extend 1.5 times the interquartile range (IQR) or the highest value above the 25^th^ and 75^th^ percentiles. The number of biological replicates and the statistical test used for each experiment are indicated in the figure legends and Methods section. No statistical method was used to predetermine sample size. No data were excluded from the analyses, and the experiments were not randomized. The investigators were not blinded to allocation during the experiments and outcome assessment.

## Data availability

All raw fastq sequencing files and processed bed files have been deposited at GEO and will publicly be available on the date of publication.

## Code availability

All code for Bullseye will be publicly available on Github (https://github.com/mflamand/Bullseye) on the date of publication.

## Ethics declarations

The authors declare no competing interests.

