## Supplemental Figures S1-S6 for "Single-molecule identification of the target RNAs of different RNA binding proteins simultaneously in cells"

**Supplemental Data**

Supplemental Fig. S1: Characterization of stable cell lines used for TRIBE-STAMP, related to Fig. 1.

Supplemental Fig. S2. Characterization of TRIBE-STAMP editing sites in HEK293T cells, related to Fig. 1.

Supplemental Fig. S3. Single-molecule co-editing of DF protein target mRNAs, related to Fig. 2.

Supplemental Fig. S4. RIP-TRIBE uncovers target mRNAs of both endogenous and overexpressed DF proteins, related to Fig. 3.

Supplemental Fig. S5. Most co-edited regions contain a single $m^6A$ site. Related to Fig. 3 and 4.

Supplemental Fig. S6. Polysome fractionation in DF-ADAR-expressing cell lines, related to Fig. 4.

Supplemental Table S1: Editing analysis from TRIBE-STAMP experiments, related to Fig. 1.

Supplemental Table S2: mRNA targets of DF protein in reanalyzed iCLIP datasets, related to Fig. 1.

Supplemental Table S3: Co-edited analysis for TRIBE-STAMP experiments, related to Fig. 2.

Supplemental Table S4: Differential editing analysis for DF-ADAR targets in DF-FLAG RIP-TRIBE, related to Fig. 3.

Supplemental Table S5: Differential editing analysis for DF-ADAR targets in endogenous DF RIP-TRIBE, related to Fig. 3.

Supplemental Table S6: Differential editing analysis for DF-ADAR targets in polysome-associated mRNAs, related to Fig. 4.

**Flamand_Fig_S1**

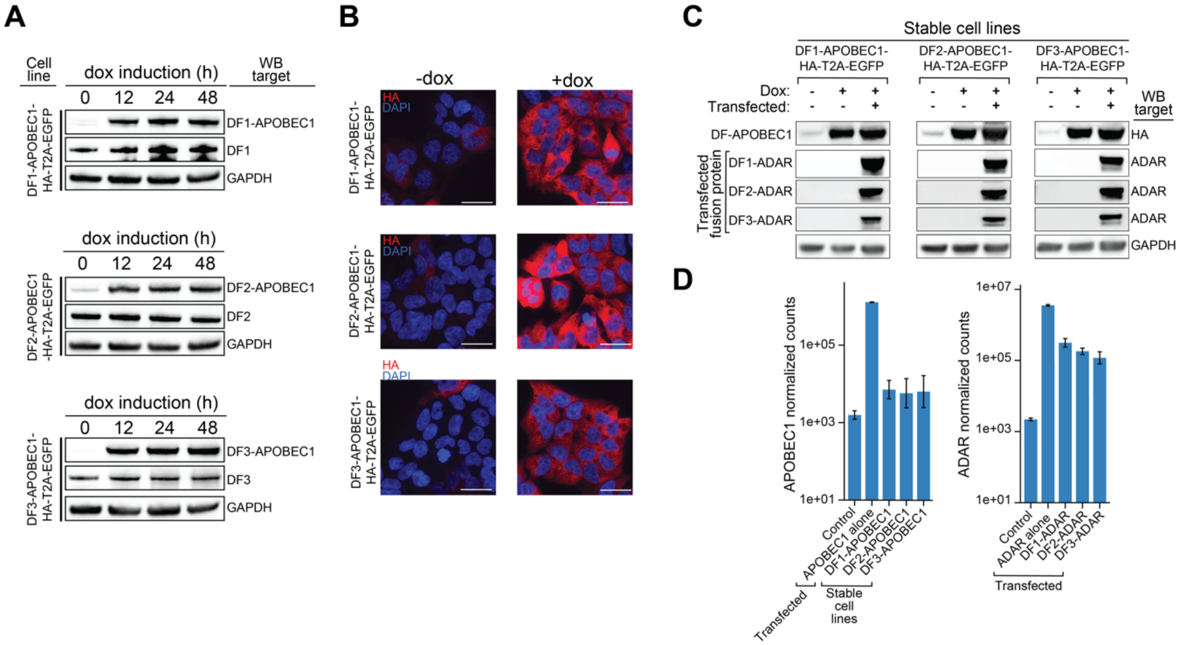


**Supplemental Fig. S1: Characterization of stable cell lines used for TRIBE-STAMP, related to Fig. 1.**

**A**, Western blot analysis of protein extracts from DF-APOBEC1-HA-T2A-EGFP stable HEK293T cell lines treated with doxycycline (dox) for 12, 24 or 48 h. Western blots were performed using antibodies for DF1, 2, or 3. The DF-APOBEC1 fusion protein products and endogenous DF protein products are indicated in separate rows. GADPH is also shown as a loading control. **B**, Immunofluorescence shows induction of DF-APOBEC1 proteins in the indicated stable HEK293T cell lines after 24 h treatment with dox. DF-APOBEC1 proteins were detected using antibodies targeting the HA epitope. Scale bar = 25 µm. **C**, Western blot analysis of protein extracts from FACS sorted EGFP/mCherry double-positive cells showing the expression of DF-ADAR and DF-APOBEC1 protein 24 h after transfection and induction with dox. Blots were probed with anti-HA and anti-ADAR antibodies. GADPH is shown as a loading control. **D**, Normalized APOBEC1 and ADAR read counts in cells subjected to TRIBE-STAMP. DF-APOBEC1-expressing stable cell lines were induced with dox for 24 h. DF-ADAR plasmids were then transfected, and cells were harvested after 24 h. Cells transfected with ADAR or APOBEC1 alone were used as controls. The mean ± standard deviation of cells expressing each fusion protein is plotted, n=9.

Flamand_Fig_S2

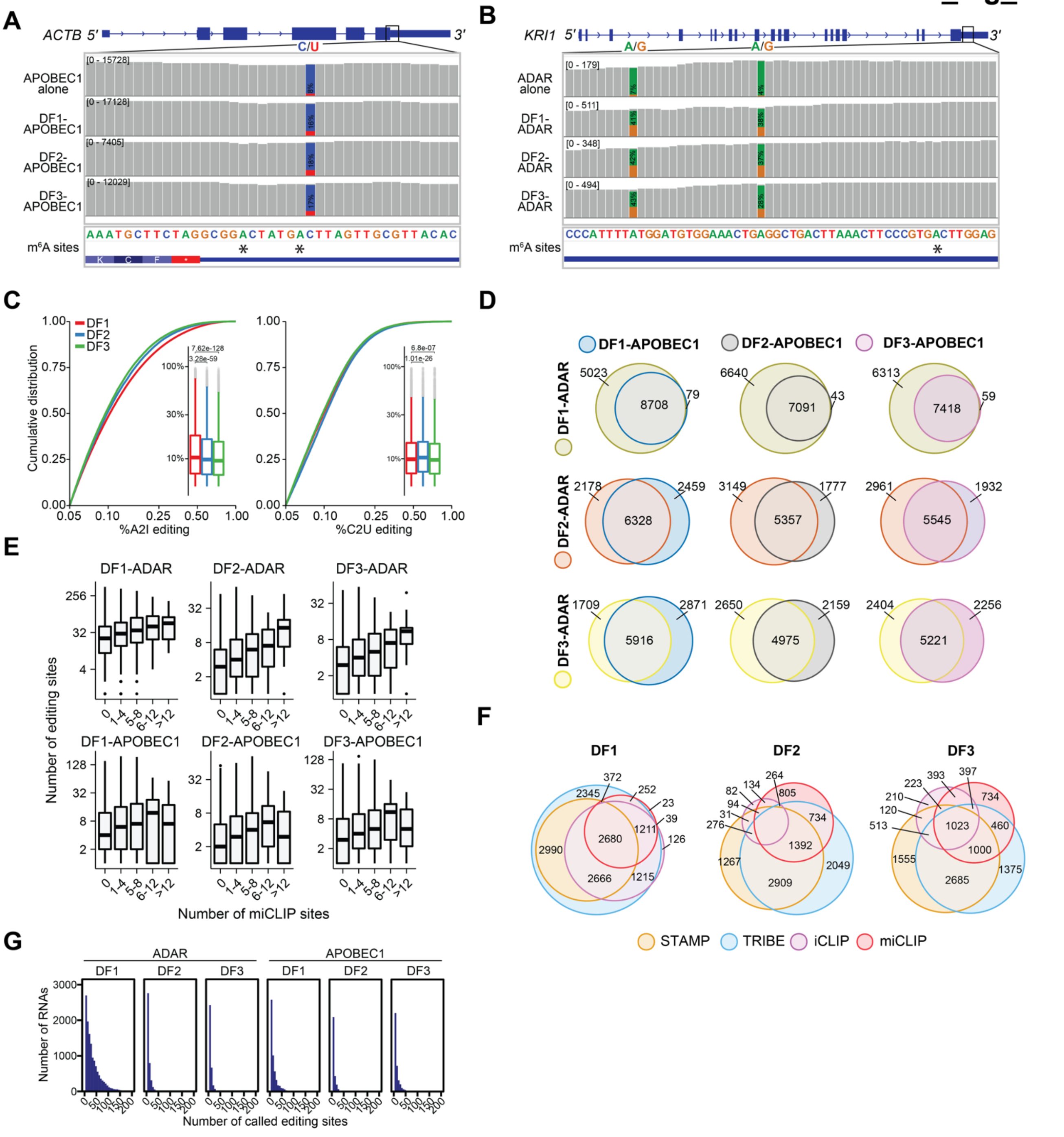

**Supplemental Fig. S2. Characterization of TRIBE-STAMP editing sites in HEK293T cells, related to Fig. 1.**

**A**, Integrative Genomics Viewer (IGV) browser read coverage tracks at the *ACTB* locus for cells expressing the indicated fusion proteins. DF-APOBEC1-expressing stable cell lines were induced with dox for 24 h. Cells transfected with APOBEC1 alone were used as control. C2U mutations found in at least 10% of reads are indicated by blue/red coloring at individual sites (blue indicates the abundance of C sites and red indicates the abundance of U sites at each position). The %C2U is indicated at called site. Each track is representative of 9 biological replicates (DF-APOBEC1) or 2 biological replicates (APOBEC1 alone). The position of mapped $m^6A$ sites by miCLIP (Linder et al. 2015) is indicated by an asterisk. **B**, Integrative Genomics Viewer (IGV) browser read coverage tracks at the *KRI1* locus for cells expressing the indicated fusion proteins. Cells were transfected with DF-ADAR or ADAR alone and RNA was harvested after 24 h. A2I mutations found in at least 10% of reads are indicated by orange/green coloring at individual sites (orange indicates the abundance of A sites and green indicates the abundance of I sites at each position, which are represented in cDNA as G). The %A2I is indicated at each called site. Each track is representative of 9 biological replicates (DF1 and DF2-ADAR), 6 biological replicates (DF3-ADAR), or 2 biological replicates (ADAR alone). The position of mapped $m^6A$ sites by miCLIP (Linder et al. 2015) is indicated by an asterisk. **C**, Cumulative distribution plots and boxplots (insets) of A2I and C2U editing rates from cells expressing the indicated DF-ADAR or DF-APOBEC1 fusion protein. *P*-values are the result of a 2-sided Wilcoxon rank sum-test. DF1-ADAR: n=450,319; DF2-ADAR: n=45,107; DF3-ADAR: n=34,713; DF1-APOBEC1: n=87,763; DF2-APOBEC1: n=32,324; DF3-APOBEC1: n=51,485. **D**, Euler plots depicting the overlap between mRNAs targeted by each combination of DF-ADAR and DF-APOBEC1 fusion proteins. **E**, Boxplots showing the number of A2I or C2U editing sites in mRNAs in relation to the number of DF iCLIP peaks in the mRNA (Patil et al. 2016). **F**, Euler plots depicting the overlap of DF target mRNAs identified by TRIBE, STAMP, or iCLIP (Patil et al. 2016) and mRNAs containing $m^6A$ sites as identified by miCLIP (Linder et al. 2015). **G**, Histogram showing the number of A2I or C2U editing sites in target mRNAs of each DF-ADAR or DF-APOBEC1 fusion protein.

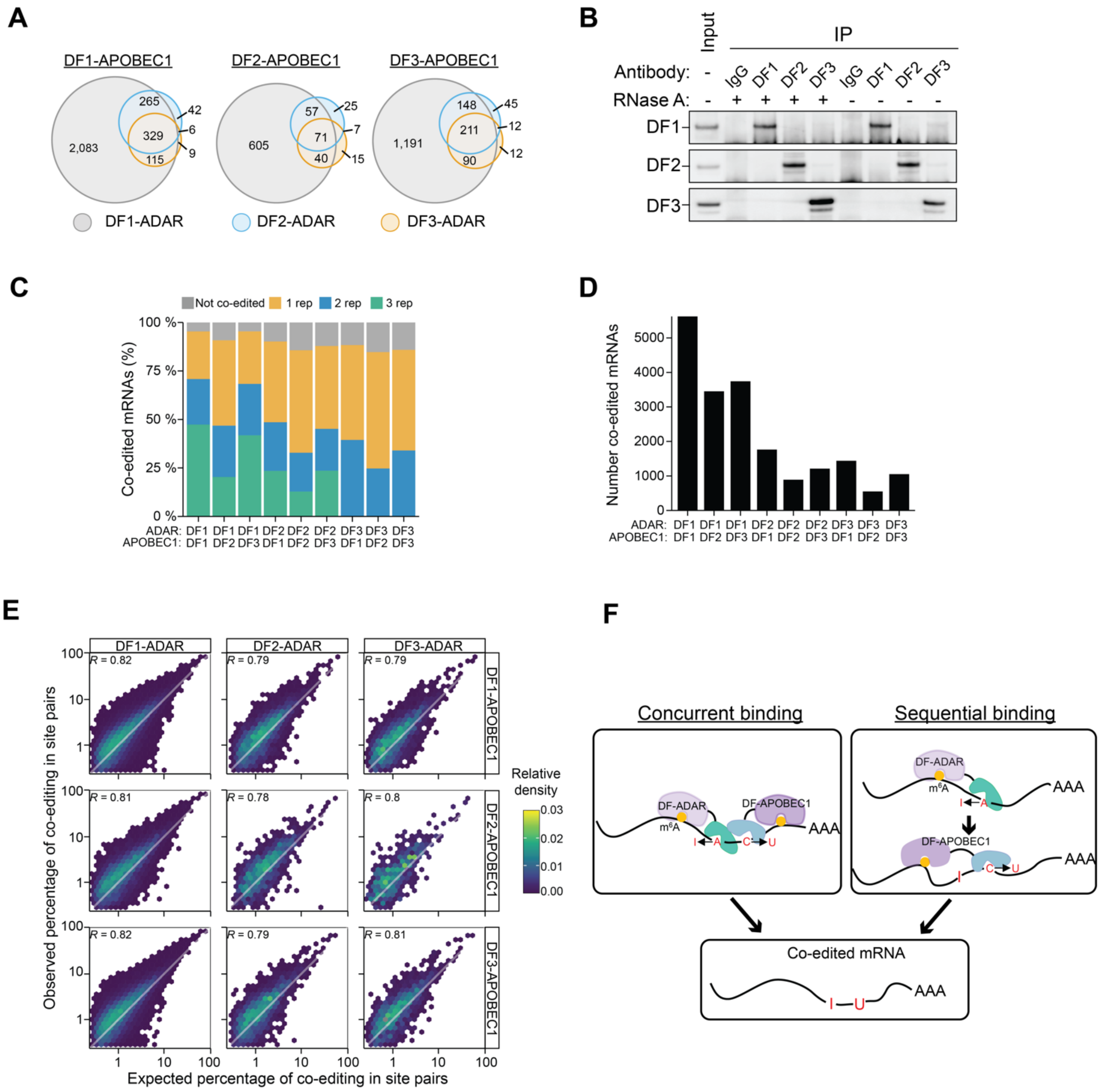


**Supplemental Fig. S3. Single-molecule co-editing of DF protein target mRNAs, related to Fig. 2 and 3.**

**A**, Overlap of co-edited mRNAs when individual DF-ADAR proteins are co-expressed with each DF-APOBEC1 protein. **B**, Endogenous DF proteins do not interact with each other in HEK293T cells. Endogenous DF proteins were immunoprecipitated in the presence and absence of RNase A, followed by western blot for each DF protein. Immunoprecipitation using IgG is shown as a negative control. Blots were probed with specific antibodies for DF1, 2 or 3. **C**, Stacked bar plot showing the proportion of transcripts co-edited in 1, 2 or 3 biological replicates. DF1-ADAR and DF2-ADAR: 3 biological replicates were performed. DF3-ADAR: 2 biological replicates were performed. **D**, Number of transcripts with co-editing in at least one biological replicate for each combination of DF-ADAR and DF-APOBEC1 fusion proteins. **E**, Co-editing is more frequent than expected by chance. 2D Density plot showing the expected and observed co-editing frequencies in editing site pairs for each combination of DF fusion proteins in TRIBE-STAMP datasets. The diagonal white line represents a trend line for a perfect linear relationship. For all DF TRIBE-STAMP combinations, the density is skewed above this curve. The Pearson correlation is indicated in the top left corner. **F**, Models for DF:RNA interactions. Concurrent binding: DF proteins bind to distinct $m^6A$ sites on the same RNA molecules. Sequential binding: DF proteins bind sequentially to the same $m^6A$ sites. Both models lead to co-editing of individual RNA molecules.

Flamand_Fig_S4

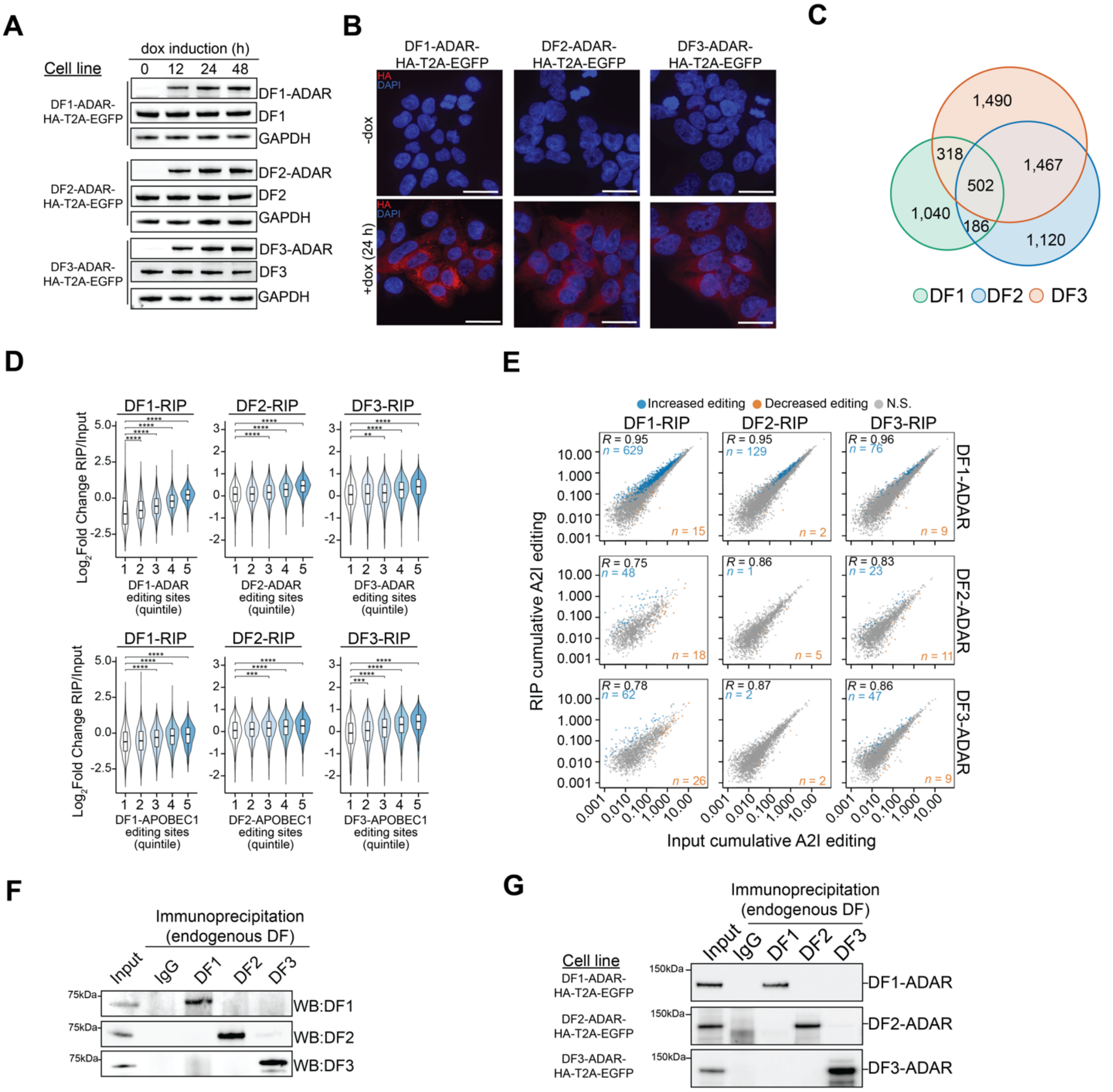

**Supplemental Fig. S4. RIP-TRIBE uncovers target mRNAs of both endogenous and overexpressed DF proteins, related to Fig. 4.**

**A**, Western blot analysis of protein extracts from DF-ADAR-HA-T2A-EGFP stable HEK293T cell lines treated with dox for 12, 24, or 48 h. Western blots were performed using DF1, 2, and 3 specific antibodies to detect both DF-ADAR fusion proteins and endogenous DF proteins. GADPH is shown as a loading control. **B**, Immunofluorescence shows induction of DF-ADAR proteins in stable DF-ADAR-HA-T2A-EGFP HEK293T cell lines treated with dox for 24 h. DF-ADAR proteins were detected using antibodies targeting the HA epitope (red) and DNA was stained with DAPI (blue). Scale bar = 25 μm. **C**, Euler diagrams showing the overlap of the mRNAs enriched in each endogenous DF RIP-seq dataset **D**, Violin plots showing the relative enrichment of mRNAs in endogenous DF RIP-seq datasets (RIP/Input). mRNAs are divided into quintiles based on the number of editing sites they contain in cells expressing the indicated DF-ADAR or DF-APOBEC1 proteins. Indicated *P-values* are the result of a Wilcoxon rank-sum test adjusted for multiple comparisons. **: ≤ 0.01, ***: ≤0.001, ****: ≤ 0.0001. **E**, Scatterplot of the cumulative RNA editing scores in input and endogenous DF RIP samples for each RIP-TRIBE combination. For each comparison, the mRNAs with a statistically significant increase or decrease in editing levels are colored in blue and orange, respectively. The statistical significance was determined using a Wald-test, FDR < 0.05. The Pearson correlation coefficient is indicated in the top left corner. The numbers of mRNAs in each category are labeled in the top left and bottom right corners.  **F-G**, Endogenous DF proteins (**D**) and DF-ADAR fusion proteins (**E**) are specifically immunoprecipitated in stable HEK293T cell lines expressing DF-ADAR-HA-T2A-EGFP. Western blot analysis of each DF protein following immunoprecipitation with individual DF antibodies or an IgG negative control is shown. Blots were probed with specific antibodies for DF1, 2, or 3. Images are representative of 3 biological replicates.

Flamand_Fig_S5

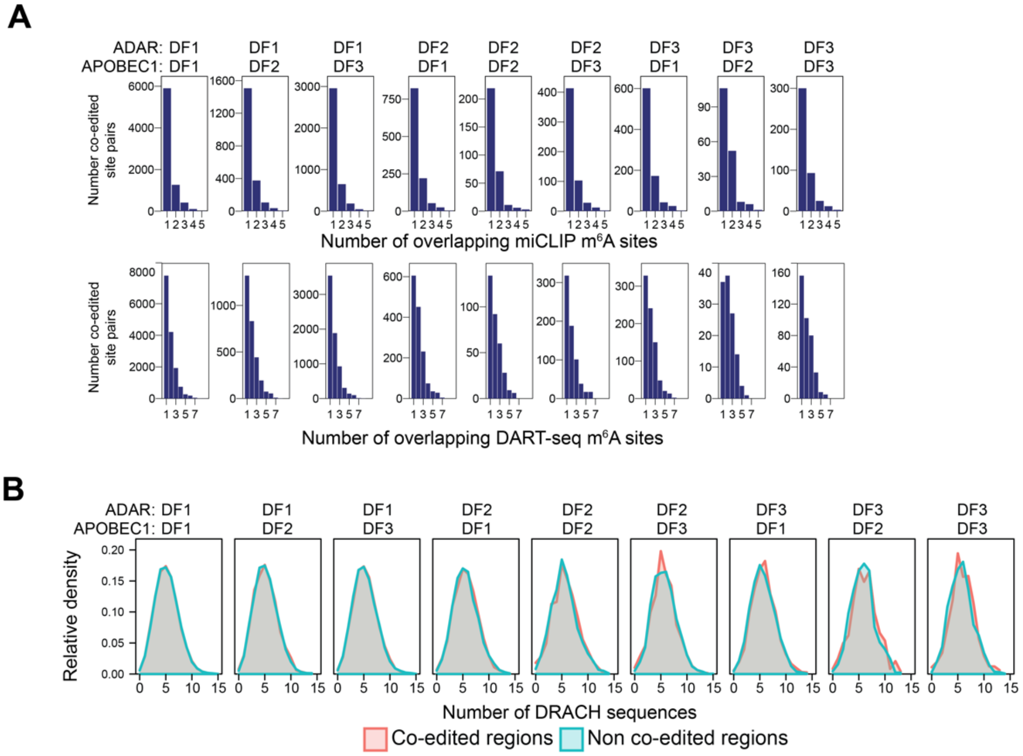


**Supplemental Fig. S5. Most co-edited regions contain a single m$^6$A site. Related to Fig. 3 and 4.**

**A**, Histograms showing the number of A2I and C2U co-edited site pairs stratified by the number of m$^6$A sites that overlap the co-edited regions. m$^6$A sites were identified by miCLIP (top row) (Linder et al. 2015) or DART-seq (bottom row) (Meyer 2019). The number of m$^6$A sites was counted in a window of fixed size (250 nt) centered on each co-edited site pair. Pairs with no overlapping m$^6$A sites are not shown. **B**, Density plots showing the number of DRACH sequences identified in fixed-size windows as in (F) for co-edited and control regions. Control regions were defined as those containing TRIBE-STAMP editing sites but lacking evidence for co-editing.

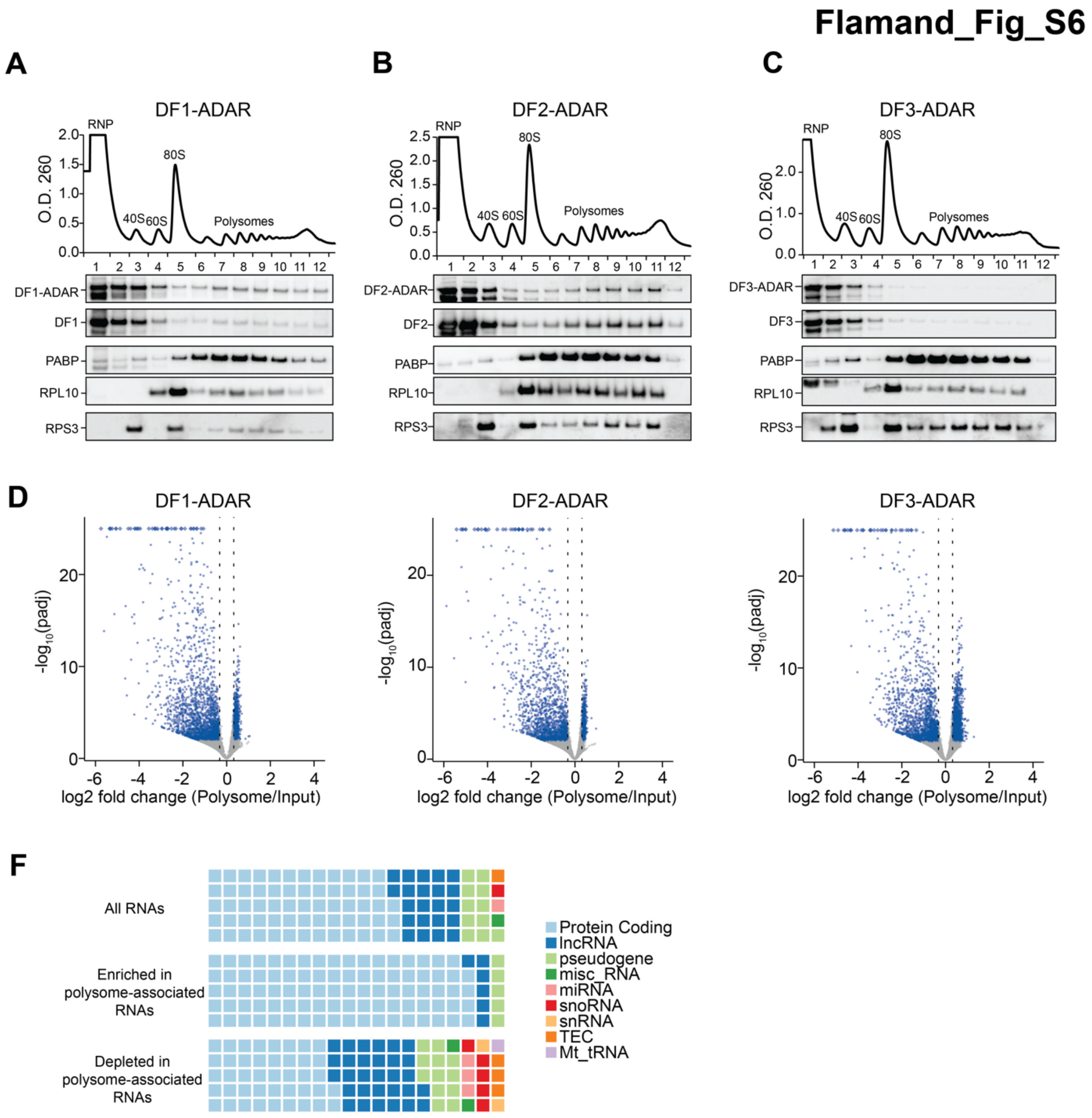


**Supplemental Fig. S6. Polysome fractionation in DF-ADAR-expressing cell lines, related to Fig. 5.**

**A-C**, Representative results of polysome fractionation for stable HEK293T cell lines expressing DF1-ADAR (a), DF2-ADAR (b), or DF3-ADAR (c). The O.D. 260 trace is shown for each cell line, in addition to western blots for individual DF proteins, DF-ADAR fusion proteins, PABP, and the ribosomal proteins RPS3 and RPL10. **D**, Volcano plots showing transcripts enriched and depleted in the polysome fractions of stable cell lines expressing the indicated DF-ADAR fusion protein. Transcripts with statistically significant enrichment or depletion from the polysome-associated RNA fraction are colored in blue (Fold Change ≥ ±1.25, FDR ≤ 0.01). **E**, RNAs enriched in the polysome fractions are overrepresented for protein coding genes. Waffle charts showing the distribution of GENCODE biotypes for all expressed RNAs and those enriched or depleted from polysomes. Each square represents 1%. TEC: to be experimentally confirmed.

**Supplemental Methods**

### *Cloning*

Coding sequences of DF proteins were cloned into pTLCV2-APOBEC1-YTH-HA-T2A-EGFP vector (Addgene plasmid # 178949) by swapping out the YTH domain to create pTLCV2-APOBEC1-DF-HA-T2A-EGFP vectors using Gibson assembly. Coding sequences of DF proteins were cloned into pcDNA3-3HA-h4E-BP1-hADARcd-E488Q vector by swapping out h4E-BP1 open reading frame. Then T2A-mCherry was fused to the C-terminal end of hADARcd-E488Q to create pcDNA3-3HA-DF-hADARcd-E488Q-T2A-mCherry vectors. These DF-hADARcd-E448Q sequences were amplified from the pcDNA3-3HA-DF-hADARcd-E488Q-T2A-mCherry vectors and then cloned into pTLCV2-APOBEC1-YTH-HA-T2A-EGFP vector by swapping out the APOBEC1-YTH to create pTLCV2-DF-hADARcd-E448Q-HA-T2A-EGFP vectors using Gibson assembly. 3xFlag was synthesized as a gene fragment (IDT) and subsequently cloned in the pCMV-APOBEC1-DF vectors by swapping out the APOBEC1 to create pCMV-3xFlag-DF vectors. psPAX2 and pMD2.g were a gift from Dr. Didier Trono (Addgene plasmid # 12260 and # 12259). pcDNA3-3HA-h4E-BP1-hADARcd-E488Q vector was a gift from Dr. Michael Rosbash.

### *Lentiviral packaging*

Lentiviral vectors pTLCV2-APOBEC1-DF-HA-T2A-EGFP or pTLCV2-DF-hADARcd-E448Q-HA-T2A-EGFP were co-transfected with psPAX2 and pMD2.g plasmids in HEK293T cells (ATCC) using Fugene HD (Promega) (Longo et al. 2013). 48 h after transfection the cell medium containing lentivirus was harvested and filtered through a 0.45 µm filter.

### *Immunofluorescence*

HEK293T stable cells were treated with 1 µg/mL of doxycycline for 24 h to induce expression of DF-hADARcd-E488Q-HA-T2A-EGFP or APOBEC1-DF-HA-T2A-EGFP.

Cells were fixed using 4% paraformaldehyde in 1X PBS for 10 min, permeabilized in 0.1% Triton X-100 in PBS for 15 min and blocked in 1% BSA in PBS for 15 min at 21°C. Cells were incubate with Rabbit anti-HA antibody for 16 h at 4 °C in 1% BSA. After 3 × 5 min washes in 1× PBS, secondary antibody (AlexaFluor488-conjugated goat anti-rabbit, 1:1,000) was then added for 1 h at 21°C. Cells were washed again in 1× PBS and incubated in 4,6-diamidino-2-phenylindole solution (1:10,000 in PBS) for 2 min. Images were acquired on a Leica DMi8 inverted fluorescence microscope.

***RNA-seq analysis***

**Alignment**

Raw sequencing reads from RNA-seq libraries were trimmed with Flexbar (v3.5.0) (Roehr et al. 2017) with the following parameters: --adapter-preset TruSeq --zip-output GZ -qf i1.8. Trimmed reads were aligned to the Ensembl GRCh38 (hg38) primary assembly, supplemented with the rat *Apobec1* sequence, using STAR (v2.7.7) (Dobin et al. 2013) in paired-end mode with the following options: --runMode alignReads --outSAMtype BAM SortedByCoordinate --readFilesCommand zcat --outSAMattributes All --outFilterMismatchNmax 20 --outFilterScoreMinOverLread 0.5 --outFilterMatchNminOverLread 0.5. Duplicate reads were then marked with STAR using the following options: --runMode inputAlignmentsFromBAM --bamRemoveDuplicatesType UniqueIdenticalNotMulti.

**Gene expression analysis**

Mapped fragments were counted to each gene in the GRCh38.102 assembly supplemented with the rat *Apobec1* gene, with featureCounts (Subread version 2.0.1) (Liao et al. 2014) and imported in R (4.1.2) for downstream processing. For the analysis of the relative expression of ADAR and APOBEC1 fusion proteins, raw counts were normalized using DESeq2 (Love et al. 2014). For differential gene expression in RIP-seq and polysome bound RNAs, DESeq2 was used to estimate fold changes. P-values were adjusted using independent hypothesis weighting (IHW) (Ignatiadis et al. 2016). For RIP-seq datasets, the size factors were computed using the type=”iterate” option.

We selected the bound RNAs for each DF RIP as those with a baseMean ≥ 25, Log2FoldChange RIP/Input ≥ 1.25 and a FDR ≤ 5%.

**(G)** Meyer KD. 2019. DART-seq: an antibody-free method for global m6A detection. *Nature Methods*.

**(H)** Patil DP, Chen CK, Pickering BF, Chow A, Jackson C, Guttman M, Jaffrey SR. 2016. m6A RNA methylation promotes XIST-mediated transcriptional repression. *Nature* **537**: 369-373.

**(I)** Roehr JT, Dieterich C, Reinert K. 2017. Flexbar 3.0 - SIMD and multicore parallelization. *Bioinformatics* **33**: 2941-2942.
